# Egocentric Spatial Scaffolds Organize Cortical Memory Engrams

**DOI:** 10.64898/2026.07.09.737550

**Authors:** Yoonsoo Yeo, Wooyoung Na, Jeehyun Kwag

## Abstract

Episodic memory requires reconstructing the position of the self within a remembered environment, yet whether and how memory engrams incorporate self-referenced spatial information remains unknown. Using activity-dependent engram tagging, longitudinal calcium imaging and optogenetic perturbation in the retrosplenial cortex, we found that cortical memory engrams are enriched for egocentric and boundary-coding neurons that encode self-position relative to environmental boundaries. Longitudinal imaging revealed that future engram neurons are preferentially recruited from a pre-existing spatial scaffold rather than generated de novo during learning. During memory retrieval, scaffold populations underwent coordinated refinement and became transiently reorganized into a scaffold–engram network state whose dynamics tracked memory expression. Silencing scaffold neurons reduced memory expression while preserving recall dynamics, whereas engram silencing abolished recall dynamics while maintaining a stable low-memory state. Together, these findings identify a scaffold–engram architecture through which episodic memories are reconstructed by reinstating the self-referenced spatial representations within remembered space.

## Introduction

Every episodic memory carries a spatial perspective — the position and orientation of the self within the environment where an experience occurred ^1-3^. This first-person perspective distinguishes episodic recollection from general semantic knowledge: to remember is not merely to know what happened, but to reconstruct where one was situated when it happened ^2,3^. How the brain represents this self-referenced spatial perspective within the cellular architecture of cortical memory remains unknown.

Memory engrams — ensembles of neurons activated during learning and reactivated during recall — provide the cellular substrate for long-term memory storage and retrieval ^4-9^. Engram studies have established which neurons are activated during memory encoding and that their reactivation is sufficient to drive recall ^4,7^. Recent work has shown that hippocampal engram neurons preferentially encode stable allocentric place fields ^10^ linking engram identity to the allocentric place cell code. However, allocentric representations — including the place cells, grid cells, and boundary representations ^11-14^— describe environmental structure independently of the observer and do not specify the remembered position of the self within that environment. Whether and how memory engrams incorporate self-referenced spatial information has not been examined.

Egocentric spatial representations have increasingly been identified across distributed cortical and hippocampal networks involved in memory and navigation, including retrosplenial, parietal, entorhinal, hippocampal, and prefrontal circuits ^15-24^. These representations encode the position and orientation of the body relative to environmental structure, providing a potential framework through which experiences may be organized relative to the self. Among these regions, the retrosplenial cortex (RSC) is particularly well positioned to mediate transformations between allocentric environmental representations and body-centered spatial frameworks ^25-29^. Computational and theoretical models have proposed that RSC integrates contextual and self-referenced information required for episodic memory reconstruction ^1,30^. Consistent with this role, RSC disruption impairs both spatial orientation and contextual memory in rodents and humans ^31-35^. Together, these findings suggest that RSC may provide a cortical substrate through which remembered experiences are reconstructed from a first-person spatial perspective.

Here we show that cortical memory engrams in RSC are not randomly organized neural ensembles but are embedded within a pre-existing egocentric spatial scaffold composed predominantly of egocentric and boundary-coding populations. This scaffold pre-exists memory formation, undergoes coordinated global refinement during recall, and becomes transiently reorganized into a coordinated network state during memory retrieval. Optogenetic dissection reveals dissociable causal contributions of scaffold and engram populations: Scaffold silencing reduced memory expression while preserving the temporal decay dynamics characteristic of contextual fear memory retrieval, indicating that the scaffold contributes to the magnitude of memory expression but does not drive retrieval dynamics. In contrast, engram silencing abolished the normal temporal evolution of recall while maintaining a stable low-memory state, indicating that the engram drives the active dynamics of memory retrieval within the scaffold framework. Together, these findings identify a scaffold–engram architecture that organizes cortical memory around self-referenced spatial representations.

## Results

### Cortical memory engrams preferentially encode self-referenced spatial representations

To determine what information cortical memory engrams encode, we performed activity-dependent engram tagging combined with longitudinal Ca^2+^ imaging in RSC during contextual fear conditioning (CFC) and recall. AAV-cFos-tTA, AAV-CaMKII-Cre, and AAV-TRE-DIO-GCaMP6s were injected into RSC to express GCaMP6s in CaMKII-positive excitatory neurons under c-Fos-tTA/TRE control (Fig. 1a). Mice underwent a three-day CFC protocol (Fig. 1b). During habituation (Day 1), mice freely explored the conditioning chamber for 30 min under the DOX-on condition, preventing activity-dependent labeling. During conditioning (Day 2), mice were transitioned to the DOX-off condition to permit c-Fos-dependent GCaMP6s expression in excitatory neurons active during memory encoding. Immediately following conditioning, mice were returned to the DOX-on condition to restrict labeling to the conditioning epoch. During recall (Day 3), mice were re-exposed to the conditioned context under the DOX-on condition, and tagged engram neurons were imaged during memory retrieval (Fig. 1b, c). Histological analysis confirmed recruitment of RSC excitatory neurons during CFC, as indicated by c-Fos expression (Extended Data Fig. 1). c-Fos-tagged GCaMP6s-expressing engram neurons were imaged during re-exposure to the conditioned context, yielding Ca²L traces from which spikes were deconvolved (Fig. 1d). Animals exhibited robust freezing during recall, confirming successful memory retrieval (Fig. 1e).

**Fig. 1.**
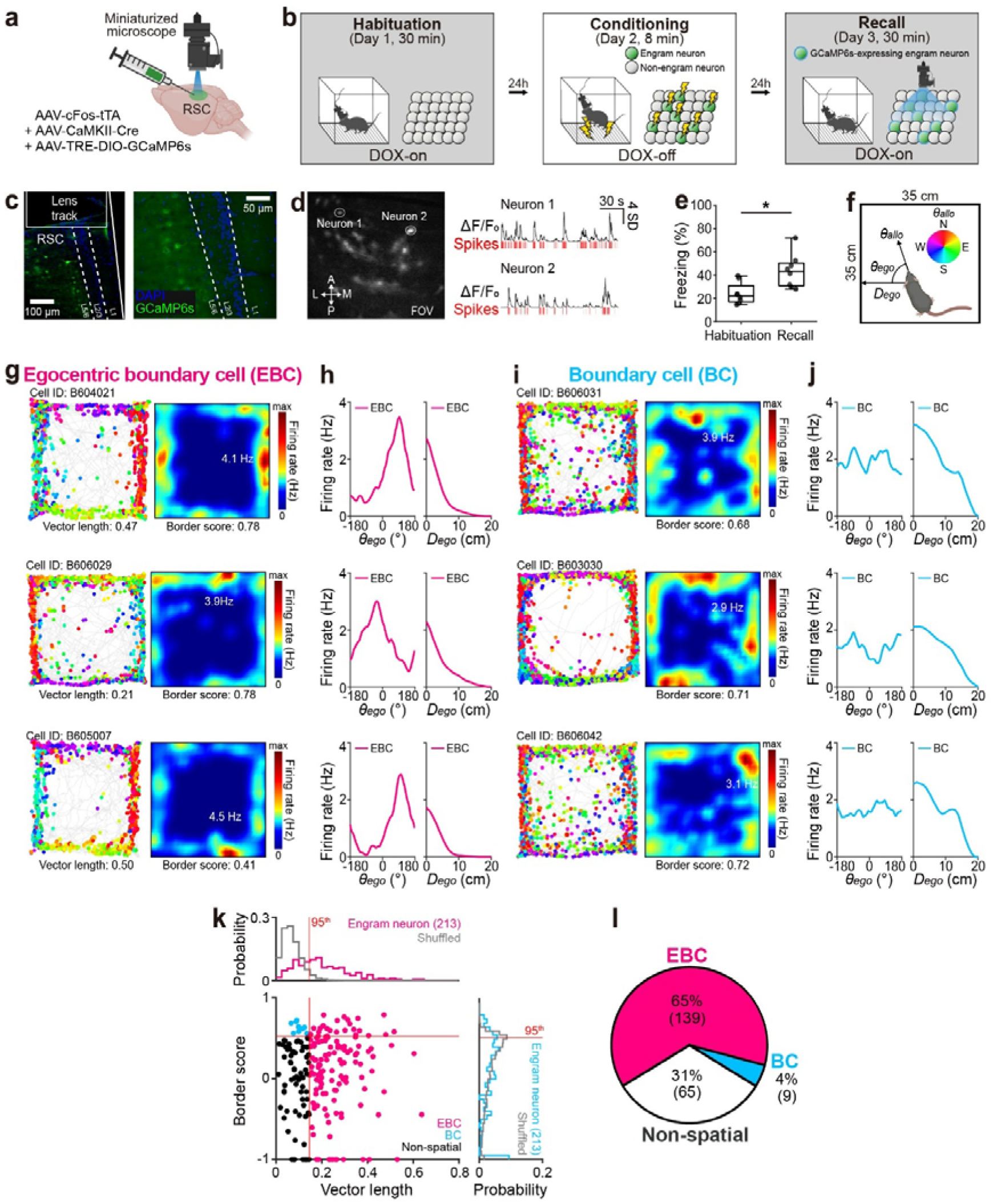
Cortical memory engrams preferentially encode the spatial position of the self. **a**, Schematic of viral injection (AAV-cFos-tTA, AAV-CaMKII-Cre, AAV-TRE-DIO-GCaMP6s), GRIN lens implantation, and Ca^2+^ imaging using miniaturized microscope of c-Fos-tagged GCaMP6s-expressing excitatory engram neurons of retrosplenial cortex (RSC) in C57BL/6J mice. **b**, Experimental timeline and doxycycline (DOX) schedule for RSC memory engram tagging during contextual fear conditioning (CFC). Mice underwent a 30-min habituation on Day 1 (DOX-on), 8-min fear conditioning with electric foot shocks for activity-dependent engram tagging on Day 2 (DOX-off), and 30-min recall for Ca^2+^ imaging of c-Fos-tagged GCaMP6s-expressing engram neurons on Day 3 (DOX-on), separated by 24-h intervals. Inset: Green filled circles: c-Fos-tagged engram neurons; Grey filled circles: non-engram neurons; Blue outline: GCaMP6s expression. **c**, Representative fluorescent image of the GRIN lens track (left) and a magnified view (right) of c-Fos-tagged GCaMP6s-expressing excitatory neurons (green) with DAPI counterstain (blue) in RSC. **d**, Representative Ca²L imaging field of view (FOV) showing c-Fos-tagged GCaMP6s-expressing neurons (left) and corresponding raw Ca²L signals (right, black traces, ΔF/F_0_) with their corresponding deconvolved spikes (red ticks) from two representative neurons. **e**, Mean freezing during habituation and recall sessions in the CFC paradigm. **f**, Illustration of the allocentric head direction (θ*allo,* color-coded for North: N, East: E, South: S, and West: W), egocentric bearing (θ*ego*), and egocentric distance (*Dego*) to the closest wall relative to the animal’s heading direction. **g**, Representative spike-trajectory plots (left; mouse trajectory: grey line, spike locations: colored dots) and the corresponding spike firing rate maps (right; color scale bar: firing rate from 0 to maximum, inset: maximum firing rate, vector length, and border score) of three representative egocentric boundary cells (EBCs) recorded from c-Fos-tagged GCaMP6s-expressing RSC engram neurons during CFC recall. Each spike is color-coded for θ*allo*. **h**, Tuning curves of θego (left) and *D*ego (right) of representative EBCs shown in (**g**). **i-j**, Same as **g-h**, but for representative non-egocentric boundary cells (BCs). **k**, Scatter plot of border score as a function of vector length for all recorded engram neurons (bottom left; dots: individual neurons), with corresponding probability distributions of vector length (top left, magenta line) and border score (bottom right, cyan line). Neurons classified as EBCs (magenta), BCs (cyan), and non-spatial neurons (black) are indicated. Grey line: randomly shuffled distributions of vector length and border score; Red line: 95^th^ percentile threshold of randomly shuffled distributions for vector length (vertical) and border score (horizontal). **l**, Pie chart showing proportions of EBCs (magenta), BCs (cyan), and non-spatial neurons (white) in c-Fos-tagged RSC engram ensembles. Box plots indicate the 25^th^, 50^th^, and 75^th^ percentiles, and whiskers represent 1.5 times the interquartile range (**e**). Statistical comparisons were performed using paired *t*-test (**e**). n = 213 neurons from 8 mice.

Spatial tuning analyses revealed that the majority of engram neurons encoded environmental boundaries relative to the self. The predominant population exhibited egocentric boundary tuning, firing selectively at specific egocentric bearing (θ*ego*), and egocentric distance (*Dego*) relative to the animal’s heading direction, and was classified as egocentric boundary cells (EBCs; Fig. 1f–h, Extended Data Fig. 2). A smaller population fired selectively proximal to environmental boundaries independent of θ*ego*, classified as non-egocentric boundary cells (BCs; Fig. 1i–j, Extended Data Fig. 2). Spatial cell type classification was performed using spike trains from the full 30-min recall session to maximize classification reliability. Classification based on egocentric vector length and border score revealed that EBCs and BCs together comprised approximately 69 % of all tagged engram neurons (Fig. 1k, l), among which 65 % were EBCs and 4 % were BCs, with neurons meeting both criteria classified as EBCs.

Together, these results demonstrate that cortical memory engrams are enriched for neurons encoding self-position relative to environmental boundaries, revealing self-referenced spatial organization as a defining principle of cortical memory engrams.

### The egocentric spatial scaffold pre-exists memory formation and undergoes global refinement during recall

To determine whether the spatial scaffold, defined as the combined population of EBCs and BCs, emerges during memory formation or pre-exists as its substrate, we performed longitudinal Ca²L imaging of GCaMP6s-expressing RSC excitatory neurons across the same three-day CFC paradigm (Fig. 2a–e). Neurons exhibiting reliable responses to foot shock during conditioning were classified as shock-responsive (SR) candidate engram neurons (40.2 %, n = 305 in 10 mice; Fig. 2f), whereas shock-non-responsive (SNR) neurons comprised the non-engram population (59.8 %, n = 453 in 10 mice; Fig. 2f).

**Fig. 2.**
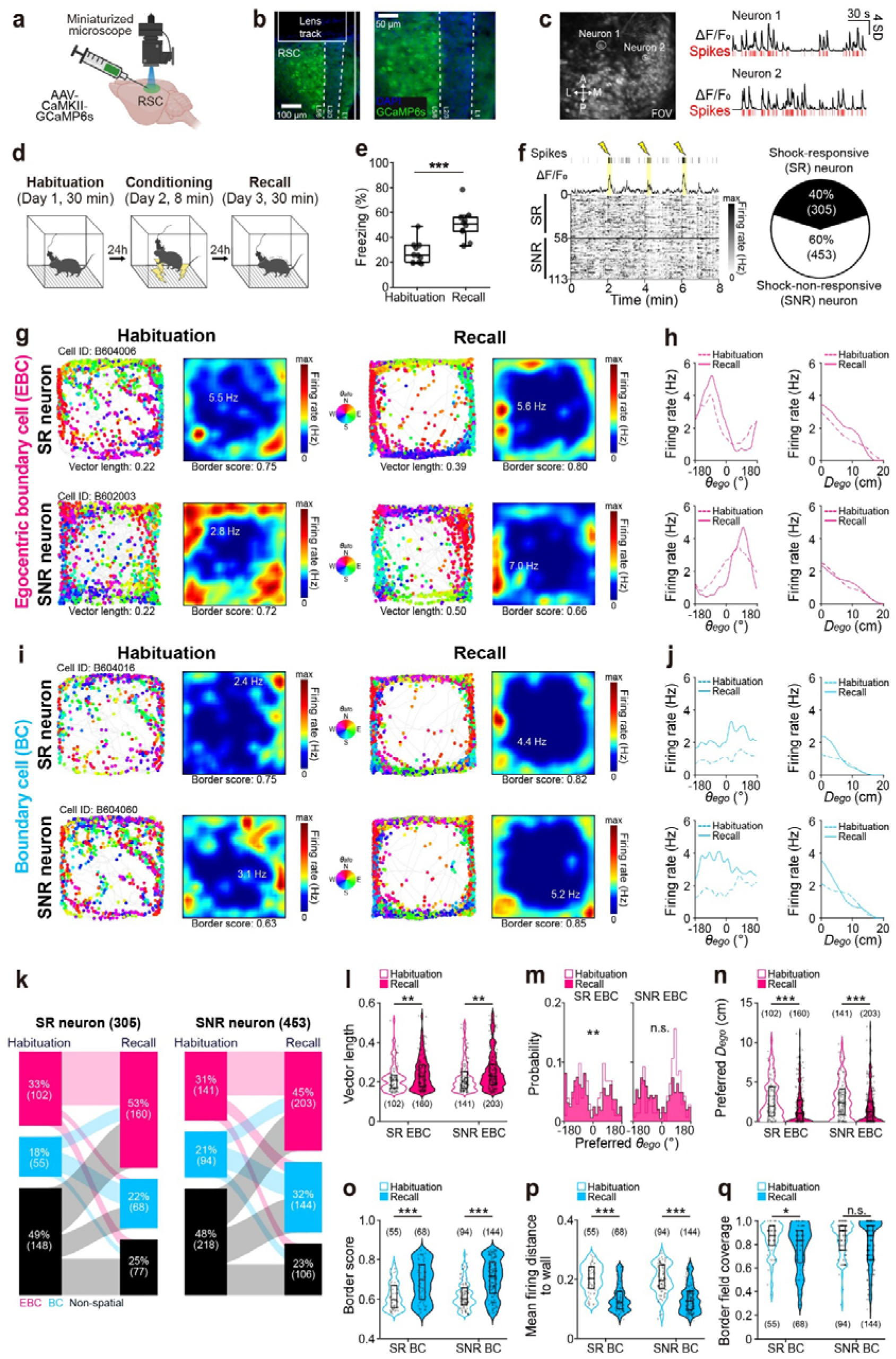
A pre-existing spatial scaffold is recruited and refined during memory formation and retrieval. **a**, Schematic of viral injection (AAV-CaMKII-GCaMP6s), GRIN lens implantation, and longitudinal Ca^2+^ imaging using miniaturized microscope of GCaMP6s-expressing neurons of retrosplenial cortex (RSC) in C57BL/6J mice. **b**, Representative fluorescent image of the GRIN lens track (left) and a magnified view (right) of GCaMP6s-expressing neurons (green) with DAPI counterstain (blue) in RSC. **c**, Representative Ca²L imaging field of view (FOV) showing GCaMP6s-expressing neurons (left) and corresponding raw Ca²L signals (right, black traces, Δ F/F_0_) with their corresponding deconvolved spikes (red ticks) from two representative neurons. **d**, Experimental timeline of the 3-day contextual fear conditioning (CFC) paradigm with longitudinal miniaturized microscope recording across all sessions. Mice underwent a 30-min habituation on Day 1, 8-min fear conditioning with electric foot shocks on Day 2, and a 30-min recall on Day 3, separated by 24-h intervals. **e**, Mean freezing during habituation and recall sessions in the CFC paradigm. **f**, (top left) Representative Ca^2+^ transients (black trace, ΔF/F_0_) and deconvolved spikes (black ticks) of a shock-responsive (SR) neuron during the conditioning session induced by electric foot shocks (yellow lightning). (bottom left) Representative firing rate maps of SR and shock-non-responsive (SNR) neurons during the conditioning session (grey scale bar: firing rate from 0 to maximum, yellow bar: delivery of electric foot shock). (right) Pie chart showing proportions of SR (black) and SNR neurons (white). **g**, Representative spike-trajectory plots (left; mouse trajectory: grey line, spike locations: colored dots) and the corresponding spike firing rate map (right; color scale bar: firing rate from 0 to maximum, inset: maximum firing rate, vector length, and border score) of SR (top) and SNR (bottom) egocentric boundary cells (EBCs) during habituation and recall sessions. Each spike is color-coded for θ*_allo_* (inset, North: N, East: E, South: S, and West: W). **h**, Tuning curves of egocentric bearing (θ_ego_, left) and egocentric distance (*D*_ego_, right) of SR EBCs (top) and SNR EBCs (bottom) recorded from GCaMP6s-expressing RSC neurons during habituation (magenta dashed line) and recall sessions (magenta line). **i-j**, Same as **g-h**, but for SR non-egocentric boundary cells (BCs) (top) and SNR BCs (bottom) during habituation (cyan dashed line) and recall sessions (cyan line). **k**, Sankey plots showing the transition of EBCs (magenta), BCs (cyan), and non-spatial neurons (black) from habituation to recall sessions for SR neurons (left, n = 305 neurons in 10 mice) and SNR neurons (right, n = 453 neurons in 10 mice). **l**, Violin plots showing the SR EBC (left) and SNR EBC vector lengths (right) during habituation (open magenta violin) and recall sessions (filled magenta violin). **m**, Probability distribution of the preferred θ*ego* of SR EBCs (left) and SNR EBCs (right) during habituation (open magenta bars) and recall sessions (fill magenta bars). **n**, Violin plots showing the preferred *Dego* of SR EBCs (left) and SNR EBCs (right) during habituation (open magenta violin) and recall sessions (filled magenta violin). **o-q**, Violin plots showing the border score (**o**), mean firing distance to wall (**p**), and border field coverage (**q**) for SR BCs (left) and SNR BCs (right) during habituation (open cyan violin) and recall sessions (filled cyan violin). Box plots indicate the 25^th^, 50^th^, and 75^th^ percentiles, and whiskers represent 1.5 times the interquartile range (**e**). In violin plots, the embedded box plots indicate the 25^th^, 50^th^, and 75^th^ percentiles (**l-q**). Statistical comparisons were performed using paired *t*-test (**e**), Watson–Williams test (**m**), and Mann–Whitney tests (**l, n-q**). n.s. denotes not significant; * *p* < 0.05, ** *p* < 0.01, *** *p* < 0.001. n = 758 neurons from 10 mice.

Using this classification, we retrospectively examined the spatial tuning of SR and SNR populations during habituation on Day 1 prior to memory formation and prospectively compared their tuning during memory recall on Day 3 (Fig. 2g–j). Representative SR and SNR EBCs exhibited selective firing at preferred θ*ego* and *Dego* during habituation that became sharper during recall, reflected by increased peak firing rates and increased boundary-proximal firing (Fig. 2g, h, Extended Data Fig. 3). Similarly, representative SR and SNR BCs exhibited stronger firing proximal to environmental boundaries during recall relative to habituation (Fig. 2i, j, Extended Data Fig. 3). These results suggest that both egocentric and contextual boundary representations are refined during memory retrieval across both engram and non-engram populations.

To quantify scaffold composition across learning, EBCs and BCs were classified using thresholds derived from habituation-session shuffle distributions (95^th^ percentile), and the same thresholds were applied to recall data to enable direct comparison across sessions. During habituation, SR and SNR populations exhibited indistinguishable spatial cell type compositions, with no significant differences in EBC or BC proportions (Fig. 2k), demonstrating that future engram neurons are recruited from the same pre-existing spatial coding population as non-engram neurons. During recall, the proportions of both EBCs and BCs increased significantly relative to habituation in SR and SNR populations alike (Fig. 2k), indicating that increased recruitment of scaffold populations occurs globally across the network rather than selectively within engram neurons.

To assess within-neuron tuning refinement, egocentric tuning and boundary coding properties were quantified for habituation-classified neurons during recall using the same habituation-derived thresholds. Vector length increased significantly from habituation to recall in both SR and SNR EBC populations (Fig. 2l), accompanied by significantly more dispersed θ*ego* distribution in SR EBCs (Fig. 2m), and reduced preferred *Dego* to environmental boundaries during recall relative to habituation (Fig. 2n). Similarly, BC populations exhibited increased border scores (Fig. 2o), decreased mean firing distance to wall (Fig. 2p), and increased border field coverage (Fig. 2q).

These effects were comparable across SR and SNR populations for vector length, preferred *Dego,* border score, and mean firing distance (Fig. 2l, n–p), while diversity of preferred θ*ego* tuning and border field coverage showed significant changes only in SR populations (Fig. 2m, q), indicating that tuning refinement is a scaffold-wide property while expansion of spatial coverage is selectively enriched in SR engram populations. When the same classification and tuning analyses were repeated using session-specific shuffle thresholds derived independently for habituation and recall, equivalent scaffold expansion and tuning refinement were observed (Extended Data Fig. 4), confirming that these findings are robust to classification threshold choice.

Together, these findings demonstrate that the egocentric spatial scaffold pre-exists memory formation and undergoes coordinated global refinement during memory retrieval across both engram and non-engram populations.

### Memory retrieval transiently reorganizes scaffold–engram network dynamics

Having established that RSC engrams are recruited from a pre-existing spatial scaffold that is refined during recall, we next asked whether memory retrieval reorganizes functional coupling within this scaffold. To address this, we analyzed pairwise correlations and population-level memory-state encoding across habituation and recall, grouping neurons into EBCs, non-egocentric BCs, and non-spatial populations. Correlations were computed within matched successive 10-min bins across habituation and recall sessions to capture temporal dynamics relative to memory expression.

Freezing declined progressively across successive recall bins, confirming that memory expression was maximal during early recall (Fig. 3a). Spike rasters of representative SR BC and SR EBC neurons revealed sparse uncoordinated firing during habituation across all bins (Fig. 3b). During recall, both populations exhibited increased and temporally coordinated activity that was strongest during early recall and progressively diminished across the later bins, paralleling the decline in freezing behavior (Fig. 3c). These patterns suggest that scaffold coordination dynamically tracks memory expression.

**Fig. 3.**
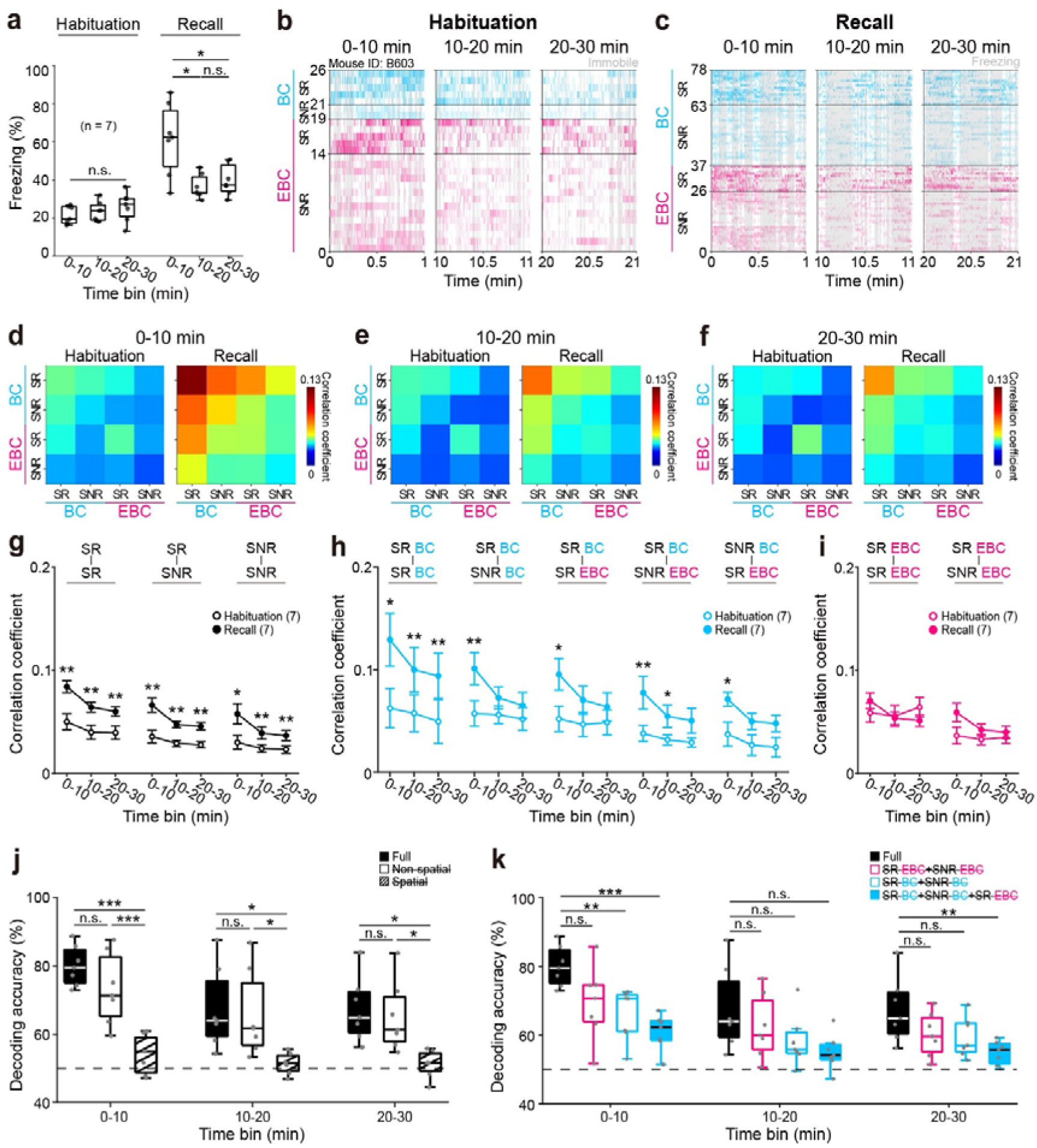
Spatial scaffold–engram coupling emerges during memory recall and tracks memory state. **a**, Mean freezing during 10-min bins of habituation and recall sessions in the contextual fear conditioning (CFC) paradigm. **b-c**, Representative spike rasters of shock-responsive (SR) and shock-non-responsive (SNR) egocentric boundary cells (EBCs, magenta) and non-egocentric boundary cells (BCs, cyan) across first 1 min of 10-min bins during habituation (**b**) and recall sessions (**c**). **d-f**, Mean pairwise correlation matrices of SR and SNR BCs and EBCs during the 0–10 min (**d**), 10–20 min (**e**), and 20–30 min time bins (**f**) across 7 mice. **g**, Mean pairwise correlations between SR-SR, SR-SNR, and SNR-SNR neuronal pairs across matched 10-min time bins during habituation (empty black circles) and recall sessions (filled black circles). **h**, Mean pairwise correlations between SR BCs or SNR BCs and other spatially tuned neuronal types (SR BC, SNR BC, SR EBC, and SNR EBC) across matched 10-min time bins during habituation (empty cyan circles) and recall sessions (filled cyan circles). **i**, Mean pairwise correlations between SR EBCs and other egocentrically tuned neurons (SR EBCs and SNR EBCs) across matched 10-min time bins during habituation (empty magenta circles) and recall sessions (filled magenta circles). **j**, Decoding accuracy across 10-min bins following ablation of spatial neurons (open), and non-spatial neurons (hatched), compared with all neurons (Full, filled). Grey horizontal dashed line: chance level (50%). **k**, Decoding accuracy across 10-min bins following ablation of SR EBC + SNR EBC (open magenta), SR BC + SNR BC (open cyan), or SR BC + SNR BC + SR EBC (filled cyan), compared with all neurons (Full, filled black). Grey horizontal dashed line: chance level (50%). Box plots indicate the 25^th^, 50^th^, and 75^th^ percentiles, and whiskers represent 1.5 times the interquartile range (**a, j, k**). Data are presented as mean ± SEM (**g-i**). Statistical comparisons were performed using repeated-measures ANOVA with post hoc Tukey’s test (**a, g-i**) and one-way ANOVA with post hoc Tukey’s test (**j-k**). n.s. denotes not significant; * *p* < 0.05, \*\**p* < 0.01, *** *p* < 0.001. n = 636 neurons from 7 mice.

Pairwise correlation matrices revealed the emergence of a structured hierarchy of functional coupling during recall that was absent during habituation (Fig. 3d–f). Boundary-coding populations exhibited the strongest coupling, egocentric populations exhibited weaker but elevated correlations, and interactions between boundary and egocentric populations showed intermediate coordination. Such hierarchical organization emerged during the first recall bin and progressively weakened across subsequent recall bins, paralleling the temporal dynamics of memory expression (Fig. 3d–f). Consistent with previous reports that memory retrieval recruits coordinated activity across engram and non-engram cortical populations^36,37^, SR-SNR coupling increased significantly during the first two recall bins relative to matched habituation bins. SR–SR and SNR–SNR correlations also increased significantly during this period, while all correlations declined during the final recall bin (Fig. 3g). At the cell-type level, all pairwise correlations involving BC populations showed significant increases during early recall bins – including SR BC–SR BC, SR BC–SNR BC, SR BC–SR EBC, SR BC–SNR EBC, and SNR BC–SR EBC pairs (Fig. 3h) – indicating that boundary-coding populations showed preferential coupling increases during recall. In contrast, EBC-EBC correlations – both SR EBC–SR EBC and SR EBC–SNR EBC pairs – showed no significant changes across any recall bin (Fig. 3i). Habituation correlations remained low and stable across all bins, confirming that coordinated scaffold dynamics emerge specifically during memory retrieval. These findings were robust to speed-based spike filtering (Extended Data Fig. 5). The comparatively lower EBC–EBC synchrony suggests that egocentric representations remain distributed across diverse self-referenced spatial configurations, whereas BC populations exhibit more globally coordinated activity during recall. One possible interpretation is that BC populations provide a stable environmental reference framework to which distributed egocentric representations are anchored during memory retrieval.

To determine whether coordinated scaffold dynamics contribute to memory-state representation, we trained a linear support vector machine (SVM) to classify freezing versus exploration from RSC ensemble activity across successive 10-min recall bins. Decoding accuracy using the full population was significantly above chance during recall (Fig. 3j). However, ablation of spatial populations (EBC + BC) reduced decoding accuracy significantly relative to the full model, while non-spatial ablation had no significant effect (Fig. 3j), indicating that memory-state information is preferentially carried by scaffold populations.

At the cell-type level, ablation of SR EBC + SNR EBC populations produced no significant reduction in decoding accuracy relative to the full model across any recall bin (Fig. 3k). Ablation of SR BC + SNR BC populations produced a significant reduction in decoding accuracy during the first recall bin (Fig. 3k). Combined ablation of SR BC + SNR BC + SR EBC populations further reduced decoding accuracy during the first recall bin and final recall bin — significantly lower than both the full model and BC-only ablation (Fig. 3k) — demonstrating that boundary-coding and egocentric populations make complementary and non-redundant contributions to memory-state decoding.

Together, these findings demonstrate that memory retrieval transiently reorganizes the spatial scaffold into a coordinated network state whose dynamics track memory expression across the recall session. Boundary-coding and egocentric populations make complementary contributions to memory-state decoding, jointly providing a functional substrate through which self-referenced spatial information is represented within the remembered context during memory retrieval.

### Dissociable causal contributions of the spatial scaffold and embedded engram to memory recall

Memory retrieval transiently reorganizes the spatial scaffold into a coordinated network state whose dynamics track expression across recall. We therefore asked whether the spatial scaffold and its embedded engram make dissociable contributions to memory retrieval by selectively silencing each population during recall using activity-dependent tagging and optogenetic inhibition.

AAV-cFos-tTA, AAV-CaMKII-Cre, and AAV-TRE-FLEX-eNpHR-EYFP were injected into RSC to express halorhodopsin under c-Fos–tTA/TRE control while control animals received AAV-TRE-DIO-EYFP without eNpHR (Fig. 4a). For spatial scaffold tagging, DOX was withheld during habituation, restricting opsin expression to habituation-active neurons (Fig. 4b). Ca^2+^ imaging of habituation-tagged GCaMP6s-expressing neurons in a separate cohort confirmed enrichment of spatially tuned populations within the tagged ensemble (Extended Data Fig. 6). For engram tagging, DOX was withheld during conditioning, restricting opsin expression to conditioning-active neurons (Fig. 4c). Histological analysis confirmed spatially restricted eNpHR-EYFP expression in RSC excitatory neurons (Fig. 4d, e).

**Fig. 4.**
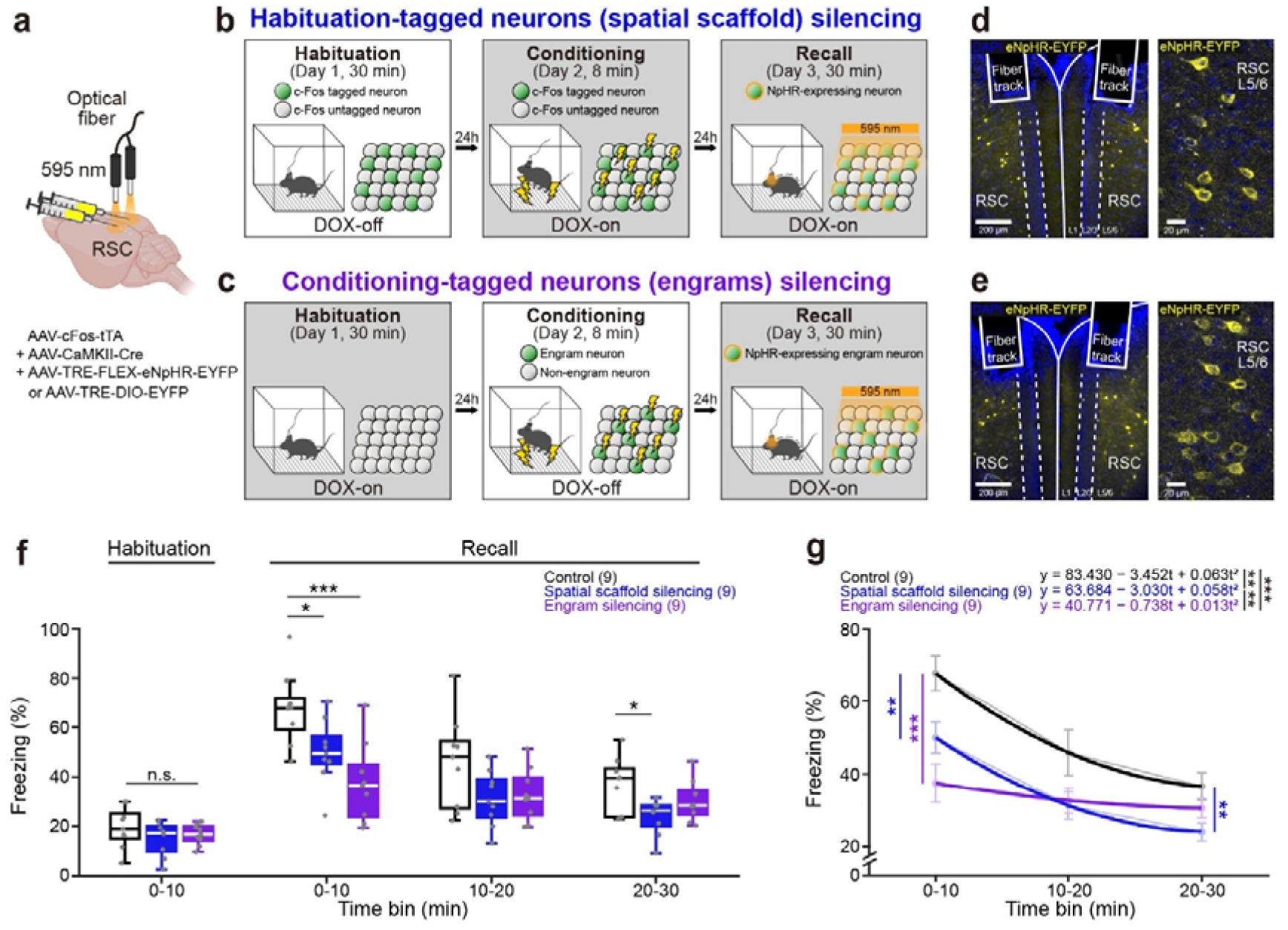
Dissociable roles of the RSC spatial scaffold and engram in memory retrieval. **a**, Schematic of viral injection (AAV-cFos-tTA, AAV-CaMKII-Cre, AAV-TRE-FLEX-eNpHR-EYFP, or AAV-TRE-DIO-EYFP), bilateral optical fiber implantation, and optogenetic inhibition of c-Fos-tagged NpHR-expressing excitatory neurons in retrosplenial cortex (RSC) in C57BL/6J mice. **b**, Experimental timeline and doxycycline (DOX) schedule for habituation activity-dependent c-Fos-tagging (spatial scaffold tagging). Mice underwent a 30-min habituation on Day 1 (DOX-off) for activity-dependent c-Fos-tagging, 8-min fear conditioning with electric foot shocks on Day 2 (DOX-on), and a 30-min recall for optogenetic silencing of spatial scaffold neurons with 595 nm light delivery on Day 3 (DOX-on), separated by 24-h intervals. Inset: Green filled circles: c-Fos-tagged neurons; Grey filled circles: c-Fos untagged neurons; Orange outline: NpHR expression. **c**, Experimental timeline and doxycycline (DOX) schedule for conditioning activity-dependent c-Fos-tagging (engram tagging). Mice underwent a 30-min habituation on Day 1 (DOX-on), 8-min fear conditioning with electric foot shocks for activity-dependent c-Fos-tagging on Day 2 (DOX-off), and a 30-min recall session for optogenetic silencing of engram neurons with 595 nm light delivery on Day 3 (DOX-on), separated by 24-h intervals. Inset: Green filled circles: c-Fos-tagged engram neurons; Grey filled circles: non-engram neurons; Orange outline: NpHR expression. **d**, Representative fluorescent images of bilateral optical fiber tracks (left) and magnified view (right) of c-Fos-tagged NpHR-expressing spatial scaffold neurons (yellow) with DAPI counterstain (blue) in RSC. **e**, Representative fluorescent images of bilateral optical fiber tracks (left) and magnified view (right) of c-Fos-tagged NpHR-expressing engram neurons (yellow) with DAPI counterstain (blue) in RSC. **f**, Mean freezing during habituation (first 10 min) and recall (three 10-min bins) session of CFC in control (empty), spatial scaffold silencing (blue), and engram silencing groups (purple). **g**, Linear mixed-effects model fit of freezing percentage over the 30-min recall session for control (black), spatial scaffold silencing (blue), and engram silencing groups (purple). Inset: quadratic model equations for each group. Box plots indicate the 25^th^, 50^th^, and 75^th^ percentiles, and whiskers represent 1.5 times the interquartile range (**f**). Statistical comparisons between groups were performed using one-way ANOVA with post hoc Tukey’s test (**f**), pairwise joint F-test, and pairwise F-test (**g**). n.s. denotes not significant; * *p*L<L0.05, ** *p*L<L0.01, *** *p*L<L0.001. n = 9 mice per group.

Silencing either spatial scaffold- or engram-tagged populations during recall reduced initial freezing relative to EYFP controls (Fig. 4f), indicating that both populations contribute to memory expression at context entry. However, their effects on recall dynamics diverged markedly across the session. Spatial scaffold silencing produced progressive memory decay across all three recall bins, similar to control mice but at a lower overall freezing level, while engram silencing produced a stable but reduced level of freezing with no significant decline across bins (Fig. 4f, g, Extended Data Fig. 7a). Linear mixed-effects modeling confirmed significant differences in freezing trajectories across the 30-min recall session (Fig. 4g).

Wall occupancy did not differ significantly across groups (Extended Data Fig. 7b). However, locomotor speed and angular head velocity were significantly elevated in engram-silenced animals relative to controls during the first recall bin only, suggesting transient increased scanning at context entry (Extended Data Fig. 7c, d).

Together, these results reveal dissociable causal contributions of the spatial scaffold and conditioning-defined engram to memory recall. Scaffold silencing reduced memory amplitude while preserving the temporal decay dynamics characteristic of contextual fear memory retrieval, indicating that the scaffold amplifies memory expression through its coordinate framework but does not drive retrieval dynamics. In contrast, engram silencing abolished temporal decay dynamics while maintaining a stable reduced memory state, indicating that the engram drives the activity dynamics of memory retrieval. This dissociation identifies the spatial scaffold as a substrate supporting memory expression and the engram as the ensemble governing the temporal dynamics of memory retrieval.

## Discussion

Episodic memory requires reconstructing the position of the self within a remembered environment, yet how cortical circuits embed the self-referenced spatial information within memory representations remains unknown. Our results show that cortical engrams in the RSC are organized around an egocentric spatial scaffold that encodes the position of the body relative to environmental boundaries. This organization anchors memory to a representation of the self relative to environmental structure. Whereas hippocampal representations provide allocentric maps of space, our findings identify a distinct cortical framework in which memory is structured around the egocentric position of the self, providing the first-person spatial perspective that defines episodic recall.

The prevailing engram framework has established which neurons are activated during memory encoding and that their reactivation drives recall ^4,7,36,38-40^, but has not addressed how memory is structured relative to the self. Fos-expressing hippocampal neurons preferentially encode stable allocentric place fields ^10^, linking engram identity to the allocentric place cell code. However, allocentric representations describe environmental structure independently of the observer and therefore do not specify where the self was situated within the remembered environment. Our findings reveal a distinct cortical organization in which RSC engrams are embedded within a spatial scaffold composed predominantly of egocentric boundary cells and boundary-coding populations that together encode self-position relative to environmental geometry. This organization is consistent with theoretical models proposing that RSC mediates transformations between allocentric maps and body-centered reference frames^1,22,28,30^.

A central question raised by this organization is whether the spatial scaffold emerges during memory formation or pre-exists as its substrate. Our longitudinal analyses demonstrate that future engram neurons are recruited from a pre-existing population that already encodes self-position relative to environmental geometry. The scaffold therefore constitutes a coordinate framework that is recruited into memory rather than generated by it. This finding has important implications for theories of engram formation: memory encoding may selectively recruit neurons from an existing spatial scaffold rather than constructing entirely new representations de novo. Such an architecture may help explain why RSC disruption produces profound contextual memory deficits ^9,41-43^: without the scaffold, the coordinate framework within which memories are organized is disrupted from the outset.

Although the scaffold pre-exists memory formation, it does not remain static. During recall, egocentric and boundary-related representations transiently strengthened across both engram and non-engram populations. This global refinement indicates that the scaffold is dynamically modulated during memory retrieval, providing an enhanced coordinate framework during active memory expression. Importantly, this refinement was distributed across both engram and non-engram populations, indicating that scaffold sharpening is a network-level property rather than an engram-specific plasticity mechanism.

Consistent with this interpretation, memory retrieval transiently reorganized the scaffold into a coordinated network state. Spatial scaffold populations showed preferential learning-dependent coordination with engram neurons during recall, a pattern absent during habituation and therefore specific to memory retrieval. Previous studies have reported increased synchrony between engram and non-engram neurons during memory retrieval ^36,37^, but the coordination observed here was spatially structured: engram neurons preferentially interacted with egocentric and boundary-coding populations rather than with non-spatial populations. This specificity suggests that memory retrieval in RSC involves reinstatement of a structured self-referenced spatial framework rather than a global increase in network activity. Population decoding analyses further demonstrated that egocentric and boundary-coding populations together contribute non-redundant information to memory-state representation, indicating that memory retrieval depends on coordinated interactions across distributed scaffold populations. One possible interpretation of this organization is that boundary-coding populations provide a stable environmental reference framework to which egocentric self-position representations are anchored during recall. Within this framework, boundary-coding populations may provide a stable environmental reference signal, whereas egocentric populations distribute self-referenced spatial information across diverse behavioral configurations, together reconstructing the self-referenced spatial information within the remembered environment during memory retrieval.

Causal perturbations revealed dissociable contributions of the scaffold and the engram to sustained memory recall. Both scaffold and engram silencing reduced initial memory expression, indicating that coordinated activity across both populations contributes to memory retrieval at context entry. However, their effects on recall dynamics diverged across the session. Scaffold silencing produced progressive memory decay, indicating that the scaffold amplifies memory expression through its coordinate framework but does not drive temporal decay dynamics characteristic of contextual fear memory retrieval. In contrast, engram silencing abolished temporal decay dynamics while maintaining a stable reduced memory state — indicating that the engram drives the active dynamics of memory retrieval within the scaffold framework. This dissociation identifies the spatial scaffold as a stabilizing substrate for cortical memory retrieval and the engram as the ensemble that shapes the temporal dynamics. A limitation of the activity-dependent tagging approach used here is the temporal imprecision of c-Fos–based labeling, which captures neurons active over a window of several hours rather than a specific behavioral epoch. More temporally precise methods — such as calcium-activated recombinase systems — would allow finer dissection of the scaffold and engram populations and their contributions to memory retrieval. Nevertheless, the behavioral dissociation between scaffold and engram silencing, combined with the enrichment of spatially tuned populations within the habituation-tagged ensemble (Extended Data Fig. 6), provides convergent evidence that the populations targeted here correspond to the spatial scaffold and memory engram, respectively.

These findings raise important questions about how egocentric cortical engrams interact with allocentric hippocampal engrams during episodic memory retrieval. Hippocampal engrams encode contextual experience and allocentric spatial information ^10,44^, whereas RSC engrams encode the egocentric position of the self within that same environment. These systems therefore appear complementary: the hippocampus represents environmental structure through allocentric place, grid cell, and boundary representations ^11-14^, whereas the RSC prominently represents the position of the self within that structure. This functional division aligns with theoretical distinctions between allocentric and egocentric reference frames that are central to models of episodic memory ^1,3,30^.

Egocentric spatial representations are increasingly being identified across multiple brain regions classically associated with memory and navigation, including medial entorhinal cortex, hippocampus, posterior parietal cortex, and prefrontal cortex ^15-21,23,24^. Although these signals have largely been studied in the context of spatial navigation and decision-making, their presence across distributed memory-related circuits raises the possibility that self-referenced spatial organization may represent a broader principle of memory processing rather than a specialization unique to RSC. Within this framework, different memory systems may encode complementary aspects of episodic experience across distinct reference frames, with allocentric representations specifying environmental structure and egocentric representations embedding the position of the self within that structure. The scaffold–engram architecture identified here therefore may reflect a general circuit motif through which memories are reconstructed from a first-person perspective across distributed cortical networks.

These findings also have implications for memory disorders. Early RSC dysfunction in Alzheimer’s disease precedes substantial hippocampal degeneration ^45,46^ and may disrupt the spatial scaffold required for normal memory retrieval. Such disruption could impair the coordinate framework necessary for accessing otherwise intact memories, potentially contributing to the disorientation and impaired memory retrieval characteristic of early disease stages. More broadly, these findings suggest that memory impairment may arise not only from loss of the engram itself, but also from disruption of the spatial framework within which memories are organized.

In summary, cortical memory engrams are embedded within a pre-existing egocentric spatial scaffold that is refined during recall, reorganized into coordinated network dynamics during memory retrieval, and required for memory expression during recall. Together, these findings identify a principle of cortical memory organization in which past experience is reconstructed by placing the self within the spatial structure of the remembered experience.

## Acknowledgement

This work was supported by the National Research Foundation of Korea (NRF) grant RS-2024-00341894 (J.K.), the Institute of Information & Communications Technology Planning & Evaluation (IITP)-ITRC (information Technology Research Center) grant funded by the Korea government (MSIT) IITP-2025-RS-2024-00436857 (J.K.) and by the New Faculty Startup Fund from Seoul National University (J.K.).

## Author contributions

J.K. conceived and designed the experiments. Y.Y. conducted the experiments for calcium imaging and analysis of the data. W.N. performed optogenetic experiments and analyzed behavior experiments. J.K. wrote the original manuscript and Y.Y. and W. N. provided feedback on the manuscript.

## Data availability

Data for this paper are available at https://github.com/SNU-NCL/EgocentricSpatialScaffold.

## Code availability

Codes for this paper are available at https://github.com/SNU-NCL/EgocentricSpatialScaffold.

