## Supplementary Materials for "Egocentric Spatial Scaffolds Organize Cortical Memory Engrams"

**Cortical Memory Engrams**

Supplementary information includes:

Materials and Methods

Extended Data Figures 1 – 7

**Methods**

**Animals**

Adult male C57BL/6J mice (#000664, DBL, South Korea), aged 3–6 months, were used for all experiments. Animals were individually housed in a temperature- and humidity-controlled vivarium under a 12-hr light/dark cycle, with *ad libitum* access to food and water except during periods requiring a doxycycline-containing (DOX) diet (DBL, South Korea). All behavioral experiments were performed during the dark cycle. All animal experimental procedures were approved by the Institutional Animal Care and Use Committee of Seoul National University (SNU-230410-1, SNU-240426-3) and conducted in accordance with institutional guidelines.

**Viral and stereotaxic surgery for *in vivo* Ca^2+^ imaging**

For surgical procedures, mice were deeply anesthetized with 1–2 % isoflurane (2 ml/min flow rate) and positioned in a stereotaxic apparatus (51730D, Stoelting Co., USA). Core body temperature was maintained at 37 °C using a homeothermic heating pad throughout surgery. For *in vivo* Ca^2+^ imaging of activity-dependent c-Fos-tagged retrosplenial cortex (RSC) excitatory neurons, a virus mixture consisting of AAV5-CaMKII0.4-Cre-SV40 (#105558-AAV5, Addgene, USA), AAV5-cFos-tTA-pA (#66794, Addgene, USA; packaged by Institute of Basic Science (IBS), South Korea) and AAV5-TRE-DIO-GCaMP6s (#183809, Addgene, USA; packaged by IBS, South Korea) at a 1:1:1 volumetric ratio was used. For longitudinal *in vivo* Ca^2+^ imaging in RSC excitatory neurons, AAV9-CaMKII-GCaMP6s (#107790-AAV9, Addgene, USA) was administered. All viral vectors were stored at –80 °C until use. The virus mixture was injected into the RSC (500 nl per site; AP: –2.5 mm, ML: –0.3 mm, DV: –0.8, –0.6, and –0.4 mm) using a Hamilton syringe (#87930, Hamilton, USA) and stereotaxic injector (#53311, Stoelting Co., USA) at an injection rate of 50 nl/min. The syringe remained at the target coordinates for an additional 5 min post-injection to facilitate viral diffusion. Subsequently, a gradient-index (GRIN) lens (1 mm diameter, 4 mm length; #1050-004605, Inscopix, USA) was implanted into the RSC (AP: –2.5 mm, ML: –0.3 mm, DV: –0.5 mm) and secured to the skull with dental adhesive resin cement (Super Bond, Sun Medical, Japan). Animals were allowed to recover for a minimum of 2 weeks before baseplate attachment. Mice were re-anesthetized with 1–2 % isoflurane (2 ml/min flow rate) to attach a magnetic baseplate (#1050-004638, Inscopix, USA). A miniaturized fluorescence microscope (nVista, Inscopix, USA) was temporarily mounted to assess the visibility of GCaMP6s-expressing neurons within the field of view (FOV). Only animals with clearly resolved GCaMP6s-expressing neurons and stable FOVs across imaging sessions were included in the dataset. If fluorescence signals were adequately detected, the baseplate was permanently affixed to the skull using dental cement (Self Curing, Vertex, Netherlands). The same microscope and focal plane were used for every imaging session with each animal to ensure consistent FOV registration across sessions. Animals were allowed to recover for at least 1 week before the behavioral experiments.

**Viral and stereotaxic surgery for *in vivo* optogenetic experiments**

For optogenetic silencing of neurons, a virus mixture consisting of AAV5-CaMKII0.4-Cre-SV40, AAV5-cFos-tTA-pA and AAV5-TRE-FLEX-eNpHR-EYFP (#89876, Addgene, USA; packaged by IBS, South Korea) at a 1:1:1 volumetric ratio was used. The control group received the same viral mixture, in which AAV5-TRE-FLEX-eNpHR-EYFP was replaced with AAV1-TRE-DIO-EYFP (#117383-AAV1, Addgene, USA). The virus mixture was injected into the RSC (500 nl, AP: −2.5 mm, ML: −0.3 mm, DV: −0.6 mm), following the same injection protocols described *in vivo* Ca^2+^ imaging. Following the virus injection, fiber-optic cannulae (Ø1.25 mm ceramic ferrule, 200 μm core, 0.50 NA, 2.5 mm length; #R-FOC-L200C-50NA , RWD, China) were bilaterally implanted 0.2 mm above the injection site at a 10° angle and fixed to the skull using dental adhesive resin cement (Super Bond, Sun Medical, Japan). Animals were allowed to recover for a minimum of 2 weeks before the behavioral experiment. Fiber placement and NpHR expression were verified histologically *post hoc* and only animals with correct targeting were included in the analysis.

**Activity-dependent c-Fos tagging of neurons**

To restrict activity-dependent c-Fos tagging, mice were maintained under the DOX-on condition (40 mg/kg doxycycline-containing diet) for at least 48 hrs prior to virus injection. To enable activity-dependent c-Fos tagging, the mice were transitioned to the DOX-off condition (standard diet) 24 hrs prior to the specific behavioral session of interest (habituation or conditioning). Immediately following the tagging session, the DOX-on condition was re-established to prevent further transgene expression and restrict tagging to the defined behavioral epoch ^1^. Depending on the viral construct used, this strategy enabled activity-dependent expression of either GCaMP6s for calcium imaging experiments or NpHR for optogenetic silencing experiments.

**Contextual fear conditioning**

For the contextual fear conditioning (CFC) paradigm, mice underwent a 3-day protocol consisting of habituation, conditioning, and recall. Prior to the experiments, mice were handled for 3 consecutive days in a separate handling chamber to reduce handling stress. All sessions were conducted in the same room maintained under constant illumination (4 lux). The conditioning chamber was cleaned with 70 % ethanol between animals. Experiments were performed in a custom-made acrylic chamber (35 cm length × 35 cm width × 25 cm height) placed on top of a custom-made square shock grid (1 m × 1 m), specifically designed to preserve natural locomotion while enabling uniform shock delivery. The shock grid consisted of alternating 5 mm-wide square steel bars and square acrylic spacers, providing a continuous flat surface while allowing effective delivery of electrical shocks. During habituation (Day 1), mice were placed in this chamber and allowed to explore for 30 min without shock delivery. After 24 hrs (Day 2, Conditioning), mice returned to the same context for a total of 8 min, during which three electric foot shocks (1 s duration, 0.5 mA amplitude) were delivered at 2-min inter-shock intervals at 2, 4, and 6 min after session onset. On Day 3 (Recall), mice were returned to the same context for 30 min in the absence of shock delivery to assess freezing behavior during memory retrieval.

***In vivo* Ca^2+^ imaging and signal processing**

For *in vivo* Ca^2+^ imaging of GCaMP-expressing neurons, mice were briefly anesthetized (< 5 min) to attach the miniaturized fluorescence microscope to the baseplate. All mice were allowed to recover from anesthesia for at least 15 min before imaging. Ca^2+^ imaging data were recorded using the Inscopix data acquisition software (IDAS, Inscopix, USA). Behavioral recordings and Ca^2+^ imaging were synchronized with a TTL-triggering system. Specifically, a TTL pulse from EthoVision XT 16 and Noldus IO box system (Noldus, Netherlands) was transmitted to nVista (IDAS, USA) to initiate simultaneous acquisition. Ca^2+^ imaging videos were acquired at a rate of 20 frames per second with 50 ms exposure time, using consistent LED power settings for each mouse across all imaging sessions. Ca^2+^ image processing was performed using Inscopix Data Processing software (IDPS, Inscopix, USA). Videos were downsampled by a binning factor of 2, and lateral brain movement was motion corrected using the image registration method ^2^. Neuronal regions of interest were identified using the PCA-ICA algorithm with an estimated neuron diameter of 18 pixels, and candidate neurons were manually curated based on spatial morphology and Ca^2+^ transient characteristics consistent with single-neuron signals. To track the same neurons across imaging sessions, longitudinal neuron registration across Days 1–3 was performed using the Inscopix longitudinal registration algorithm. This process aligned neuron maps by rigid transformation and matched individual neurons based on their normalized cross-correlation scores ^3^. Only neurons successfully registered across all three sessions were included in longitudinal analyses. Ca^2+^ traces were normalized to the baseline standard deviation (SD) of each neuron. Spike trains were estimated from $\Delta$F/F_0_ traces using OASIS-based deconvolution with a decay time constant of 1 s ^4^.

**Optogenetic modulation of activity-dependent c-Fos-tagged RSC neurons**

For optogenetic modulation of activity-dependent c-Fos-tagged RSC excitatory neurons, DOX-on/off conditions were managed as described above. Optogenetic inhibition was achieved by delivering yellow light (595 nm, 1 mW at the fiber tip) using an LED driver (LEDD1B, Thorlabs, USA) connected to bifurcated fiber-optic cables (BFYL2LF01, Thorlabs, USA). The light power at the fiber tip was calibrated to 1 mW before each experimental session using an optical power meter (PM100D, Thorlabs, USA). Continuous illumination was delivered bilaterally throughout the entire 30-min recall session to optogenetically suppress NpHR-expressing c-Fos-tagged neuronal population. Such continuous optogenetic silencing protocol has previously been shown to reduce neuronal firing during prolonged light delivery without detectable light-induced neuronal injury responses ^5-7^.

Two distinct optogenetic-silencing experiments were conducted. For silencing habituation activity-dependent c-Fos-tagged neurons (spatial scaffold), NpHR expression was restricted to neurons active during habituation by maintaining mice under the DOX-off condition during the habituation session (Day 1). Immediately following habituation, mice were returned to the DOX-on condition to prevent further transgene expression. Yellow light was delivered through the implanted optic fiber during the recall session to inhibit scaffold neurons.

For silencing conditioning activity-dependent c-Fos-tagged neurons (engram), NpHR expression was restricted to neurons active during memory encoding by maintaining mice under DOX-off condition during the conditioning session (Day 2). Immediately following conditioning, mice were returned to the DOX-on condition to prevent further transgene expression. Yellow light was delivered through the implanted optic fiber during the recall session to inhibit engram neurons. Control animals expressing EYFP alone underwent identical DOX diet light delivery procedures with spatial scaffold silencing group.

**Behavior tracking and analysis**

Animal position and head direction were tracked from behavioral videos using DeepLabCut (DLC) ^8^. A custom DLC pose-estimation model was trained using 100 manually labeled frames selected from a representative subset of behavioral videos. Five body parts (snout, left ear, right ear, body center, and tail base) were annotated and used for pose estimation. The model was trained for 100,000 iterations and subsequently applied to the full behavioral video dataset. The tracking results were visually inspected in DLC-generated labeled videos.

To ensure adequate spatial sampling for egocentric and boundary tuning analysis and contextual fear conditioning analysis, spatial coverage of the behavioral arena was quantified for each animal. The chamber was divided into 0.5 cm × 0.5 cm spatial bins, and coverage was calculated as the fraction of spatial bins visited at least once during the session relative to the total chamber. Mice with spatial coverage below 0.59 (59 %) during the habituation session were excluded from subsequent analyses. Speed was calculated from the frame-to-frame displacement of the body center coordinates. Freezing was defined as periods during which speed remained < 0.5 cm/s for at least 0.5 s ^9, 10^. All other periods were classified as movement (unfreezing) periods. Wall occupancy was calculated as the percentage of time during which the body center was located within 5 cm of the chamber boundary. Head direction was defined as the angle of the vector extending from the midpoint of the left and right ears to the snout, relative to a fixed reference axis in the behavioral arena. Angular head velocity (AHV) was computed as the frame-to-frame changes in head direction angle.

**Border score analysis**

Boundary cells were identified through spatial firing rate map analysis and calculation of the border score (*b*), as previously established ^11^. The experimental chamber was discretized into 1.5 cm × 1.5 cm spatial bins. Raw firing rates for each bin were calculated by dividing the total number of spikes by occupancy time. Bins with an occupancy < 0.05 s were excluded from the analysis. The resulting raw rate maps were smoothed by convolution with a two-dimensional Gaussian kernel (5-bin width, $\sigma$ = 5 bins). Border fields were defined as contiguous spatial bins with firing rates $\geq$ 30 % of the peak firing rate. The border score *b* was calculated as:

$$b=\frac{c_{M}-d_{m}}{c_{M}+d_{m}}$$

where $c_{M}$ is the maximum border field coverage by any single border field, and $d_{m}$ is the mean firing distance to the nearest wall. The *b* ranges from −1 to +1, where values approaching +1 indicate a strong preference for boundaries, while values near −1 indicate border fields centered far from the walls. A neuron was characterized as a boundary cell (BC) if its border score exceeded the 95^th^ percentile of a shuffled null distribution. This null distribution was generated for each cell by performing 1,000 circular time-shifts of the spike train relative to the animal's trajectory, thereby decoupling the spikes from specific boundary locations while preserving the temporal firing structure.

**Egocentric boundary vector analysis**

Egocentric boundary representations were analyzed by calculating spike firing rates as a function of the animal’s egocentric relation to environmental boundaries ^12-15^. For each video frame, boundary distances were binned into 1.5 cm intervals, and egocentric bearing angles relative to the animal's allocentric head direction were binned from 0° to 360° in 3° bins, where 0° was defined as directly ahead, −90° to its left, ±180° directly behind, and 90° to its right. Egocentric boundary representations were analyzed only during movement epochs (speed > 2 cm/s), within bins with an occupancy > 0.05 s, and at least one spike, ensuring reliable spatial sampling. Egocentric vector was then calculated by dividing the number of spikes within each 3° × 1.5 cm bin by the total occupancy time for that bin. Raw egocentric vector rates were smoothed using a two-dimensional Gaussian kernel (5-bin width, $\sigma$ = 5 bins). To identify neurons with significant egocentric boundary vector coding, the mean resultant vector of the cell's egocentric directional firing was calculated, collapsing across all distances to the boundary. The mean resultant vector was calculated as:

$$R= \frac{1}{nm}\sum_{\theta=1}^{n} \sum_{D=1}^{m} F_{\theta,D}e^{i\theta}$$

where *θ* represents the orientation relative to the animal, *D* the distance from the animal, $F_{\theta,D}$ the firing rate in each orientation-by-distance bin, *n* the number of orientation bins, *m* the number of distance bins, *e* Euler's constant, and *i* the imaginary unit. The mean vector length (VL), defined as the absolute value of the mean resultant vector $|R|$, was used to quantify the strength of egocentric bearing tuning. A neuron was characterized as an egocentric boundary cell (EBC) if its VL exceeded the 95^th^ percentile of a shuffled null distribution. This null distribution was generated for each cell by performing 1,000 circular time-shifts of the spike train relative to the animal's trajectory, thereby decoupling the spikes from specific egocentric boundary locations while preserving the temporal firing structure.

**Shock-responsive neuron analysis**

To quantify shock-responsiveness, deconvolved spike firing rates were analyzed within a 3 s window immediately following foot shock onset (defined as the "shock response"). A neuron was classified as shock-responsive (SR) if its measured shock response significantly exceeded the 99^th^ percentile of its shuffled null distribution, generated by 1,000 circular time-shifts of the spike train relative to shock onset. Neurons not meeting this criterion were defined as shock-non-responsive (SNR).

**Synchrony analysis**

To quantify functional connectivity among neural populations, pairwise Pearson correlation coefficients were computed between individual neurons. For this analysis, only mice with a minimum of three neurons in each group across both habituation and recall sessions were included to ensure reliable estimation of pairwise correlations. Deconvolved spike data were first smoothed using a Gaussian kernel ($\sigma$= 100 ms) to obtain a continuous estimate of the neural activity for each neuron. The smoothed activity of each neuron was then normalized using a z-score transformation within each session to ensure that units with different activity levels contributed equally to the subsequent analysis. To assess temporal changes in network synchrony, Pearson correlation matrices were computed between all neuronal pairs using the smoothed population activity traces within successive 1-min bins across the full 30-minute session. The resulting correlation values were then averaged across three successive 10-min time bins (0-10, 10-20, and 20-30 min) for both habituation and recall sessions, yielding mean pairwise correlations per time bin for each animal.

**Behavior decoding**

To evaluate whether RSC population activity encoded memory-related behavioral states, a linear support vector machine (SVM) classifier was employed to decode freezing behavior from population activity during the first 10 min of the recall session ^16^. For this decoding analysis, only mice with a minimum of three neurons in each group across both habituation and recall sessions were included to ensure sufficient population size for reliable classification. Deconvolved spike data was first smoothed using a Gaussian kernel ($\sigma$= 100 ms) to obtain a continuous estimate of neural activity. The smoothed activity of each neuron was then normalized using a z-score transformation within each session to construct the feature matrix. Target labels correspond to binary behavioral states (freezing vs. unfreezing) sampled at each time bin. To ensure a 50 % theoretical chance level and account for potential label imbalances, the dataset was balanced prior to classifier training through 100 iterations of randomly under-sampling the majority behavioral label to match the size of the minority label. Classifier performance and within-session generalizability were assessed using a 10-fold cross-validation scheme within the recall session. For each cross-validation fold, a linear SVM classifier was trained on a randomly selected 90 % of the temporal bins using custom-written MATLAB (MathWorks, USA) code and decoding accuracy was quantified as the proportion of correctly classified bins within the held-out 10 % test set. To isolate the functional significance of specific neural subgroups, an ablation analysis was performed. The functional contribution of each neural subgroup was evaluated by systematically applying 100 iterations of random circular shifting to the activity of each neuron within the feature matrix prior to classifier training and testing, and subsequently comparing decoding performance to that of the full population.

**Quadratic mixed-effects analysis of freezing behavior**

Freezing dynamics across the 30-min recall session were analyzed using linear mixed-effects models with quadratic time terms ^17^, implemented using the ‘fitlme’ function in the MATLAB Statistics and Machine Learning Toolbox (MathWorks, USA). The midpoint of each recall bin (in min) and its squared term (Time²) were included as continuous fixed effects, and mouse groups (Control, spatial scaffold silencing, and engram silencing) were included as a categorical fixed effect. Group × Time and Group × Time² interaction terms were included as the primary variables of interest to assess differences in recall dynamics across groups. Animal identity was included as a random effect with mouse-specific intercepts and linear and quadratic time slopes. Pairwise differences between fitted quadratic curves were assessed using F-tests on fixed-effect contrasts, comparing intercept, linear (Time), and quadratic (Time^2^) components between groups.

**Histology and immunohistochemistry**

For histological imaging of the mouse brain, mice were deeply anesthetized with Avertin (T48402, Sigma-Aldrich, USA) and perfused with ice-cold 4 % paraformaldehyde (PFA, 158127, Sigma-Aldrich, USA), followed by fixation in 4 % PFA at 4 °C overnight, then cryoprotected in 30 % sucrose in phosphate-buffered saline (PBS) for 24-48 hrs. The tissue was embedded in optimal cutting temperature (OCT) compound (Tissue-Tek O.C.T., SAKURA, Japan) and frozen at –80 °C prior to sectioning.

Coronal brain sections (50 μm thickness) were cut using a cryostat (YD-2235, Jinhua Yidi Medical Appliance Co., China), rinsed three times for 5 min in PBS to remove residual OCT, and mounted on glass slides with antifade medium containing DAPI (Vectashield, Vector Laboratories, USA). Fluorescent signals were visualized and imaged using a confocal laser-scanning microscope (LSM980, Zeiss, Germany) and fluorescence microscope (DM2500, Leica, Germany).

To identify RSC excitatory neurons activated during CFC habituation and conditioning, c-Fos immunohistochemistry was performed. AAV9-CaMKII-GCaMP6s (#107790-AAV9, Addgene, USA) was injected into the RSC as described above, and mice were sacrificed 1 hr after the end of the habituation or conditioning session. Coronal sections were prepared as described above and incubated in blocking solution consisting of PBST (PBS with 0.3 % Triton X-100; Triton X-100, Sigma-Aldrich, USA) and 5 % normal donkey serum (#566460, Merck, USA) for 2 hrs at room temperature. Sections were subsequently incubated with primary antibody (1:500, rabbit anti-c-Fos, #2250, Cell Signaling Technology, USA) at 4 °C overnight. The following day, sections were washed three times with PBS (5 min each) and incubated with secondary antibody (1:500, #A10042, Alexa Fluor 568 donkey anti-rabbit IgG, Invitrogen, USA) for 1 hr at room temperature. After three additional washes in PBS (10 min each), sections were mounted and imaged using fluorescence microscope (DM2500, Leica, Germany).

**Statistical analysis**

All statistical analyses were conducted in MATLAB (MathWorks, USA). Paired Student’s *t* test, Mann–Whitney tests, one-way, two-way, or repeated-measures ANOVA with *post hoc* Tukey’s test, and F-test were used as appropriate. For tuning comparisons, permutation tests were used. For circular data, Watson–Williams test was used. Significance was set at *p* < 0.05. Data are reported as mean ± SEM.

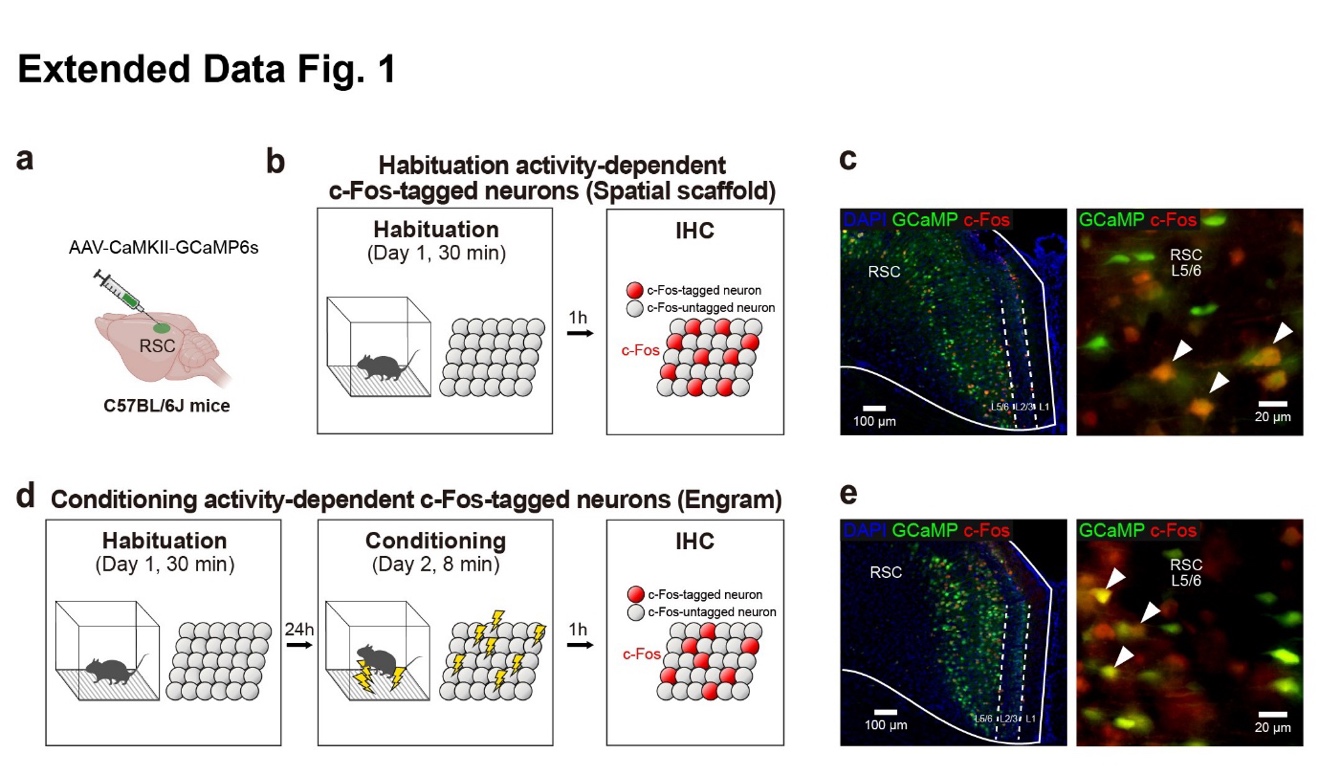

**Extended Data Fig. 1 Validation of activity-dependent c-Fos tagging in RSC excitatory neurons.**

**a**, Schematic of viral injection (AAV-CaMKII-GCaMP6s) to express GCaMP6s in excitatory neurons of retrosplenial cortex (RSC) in C57BL/6J mice.

**b**, Experimental timeline for detecting habituation-expressed c-Fos. Mice underwent a 30-min habituation on Day 1, followed by immunohistochemistry (IHC) 1 h later. Inset: Red filled circles: c-Fos-tagged neurons; Grey filled circles c-Fos-untagged neurons.

**c**, Representative fluorescent images (left) and a magnified view (right) showing the expression of GCaMP6s (green) and c-Fos (red) with DAPI counterstain (blue) in the RSC. Arrowheads: GCaMP6s and c-Fos co-expressing neurons.

**d**, Experimental timeline for detecting conditioning-expressed c-Fos. Mice underwent a 30-min habituation on Day 1, 8-min conditioning receiving electric foot shocks on Day 2 after a 24-h interval, followed by IHC 1 h later. Inset: Red filled circles: c-Fos-tagged neurons; Grey filled circles: c-Fos-untagged neurons.

**e**, Representative fluorescent images (left) and a magnified view (right) showing the expression of GCaMP6s (green) and c-Fos (red) with DAPI counterstain (blue) in the RSC. Arrowheads: GCaMP6s and c-Fos co-expressing neurons.

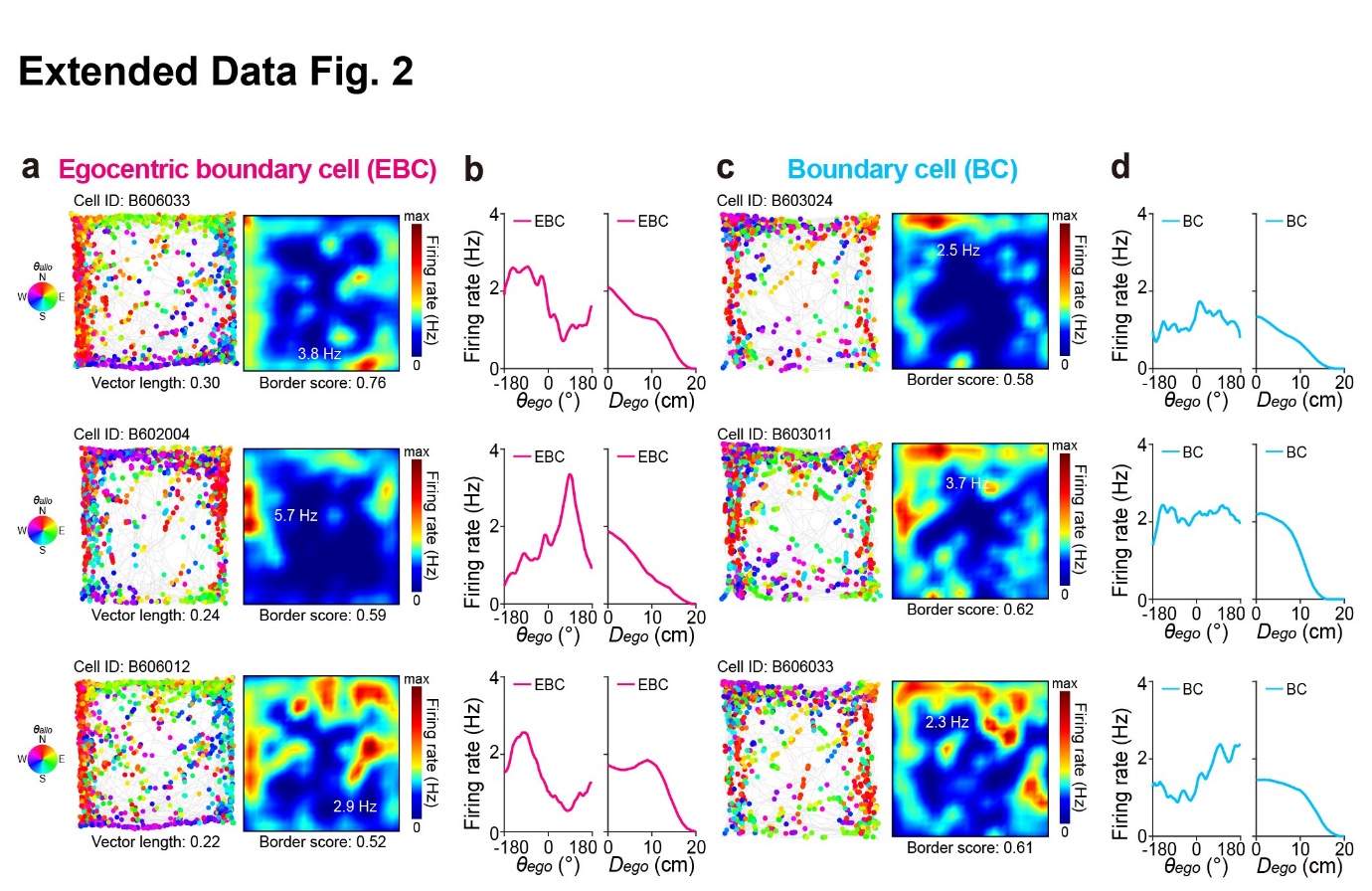

**Extended Data Fig. 2 Representative spatial tuning of egocentric boundary cells and boundary cells in RSC memory engram neurons.**

**a**, Representative spike-trajectory plots (left; mouse trajectory: grey line, spike locations: colored dots) and the corresponding spike firing rate maps (right; color scale bar: firing rate from 0 to maximum, inset: maximum firing rate, vector length, and border score) of three representative egocentric boundary cells (EBCs) recorded from c-Fos-tagged GCaMP6s-expressing RSC engram neurons during CFC recall . Each spike is color-coded for *θallo* (inset, North: N, East: E, South: S, and West: W).

**b**, Tuning curves of egocentric bearing (*θ*ego, left) and egocentric distance (*D*ego, right) of EBCs shown in (**a**).

**c-d**, Same as **a-b**, but for representative non-egocentric boundary cells (BCs).

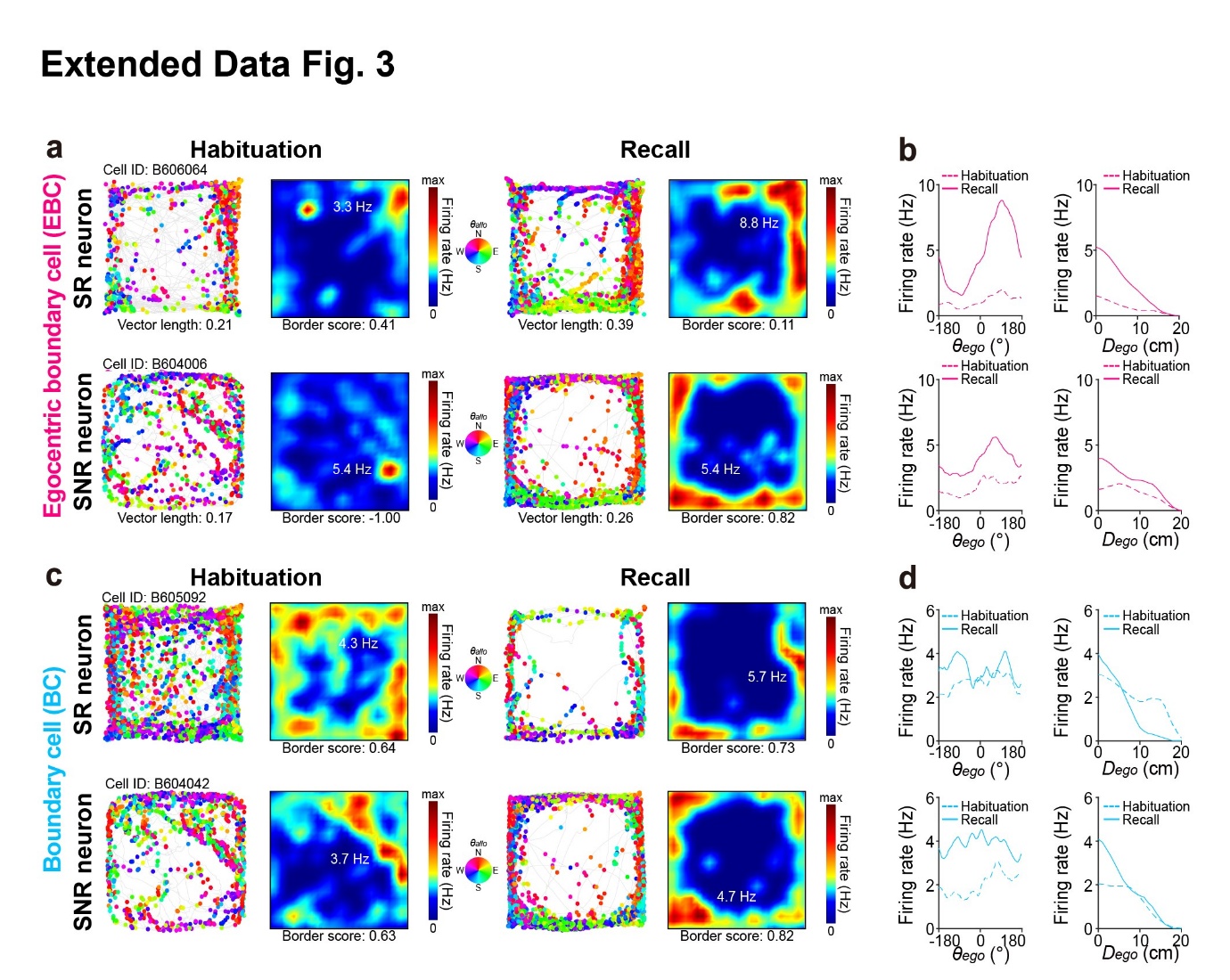

**Extended Data Fig. 3 Representative spatial scaffold refinement from habituation to recall in RSC engram and non-engram neurons.**

**a**, Representative spike-trajectory plots (left; mouse trajectory: grey line, spike locations: colored dots) and the corresponding spike firing rate maps (right; color scale bar: firing rate from 0 to maximum, inset: maximum firing rate, vector length, and border score) of SR (top) and SNR (bottom) egocentric boundary cells (EBCs) during habituation and recall sessions. Each spike is color-coded for *θallo* (inset, North: N, East: E, South: S, and West: W).

**b**, Tuning curves of egocentric bearing (*θ*ego, left) and egocentric distance (*D*ego, right) of SR EBCs (top) and SNR EBCs (bottom) recorded from GCaMP6s-expressing retrosplenial cortex neurons during habituation (magenta dashed line) and recall sessions (magenta line).

**c-d**, Same as **a-b**, but for SR non-egocentric boundary cells (BCs) (top) and SNR BCs (bottom) during habituation (cyan dashed line) and recall sessions (cyan line).

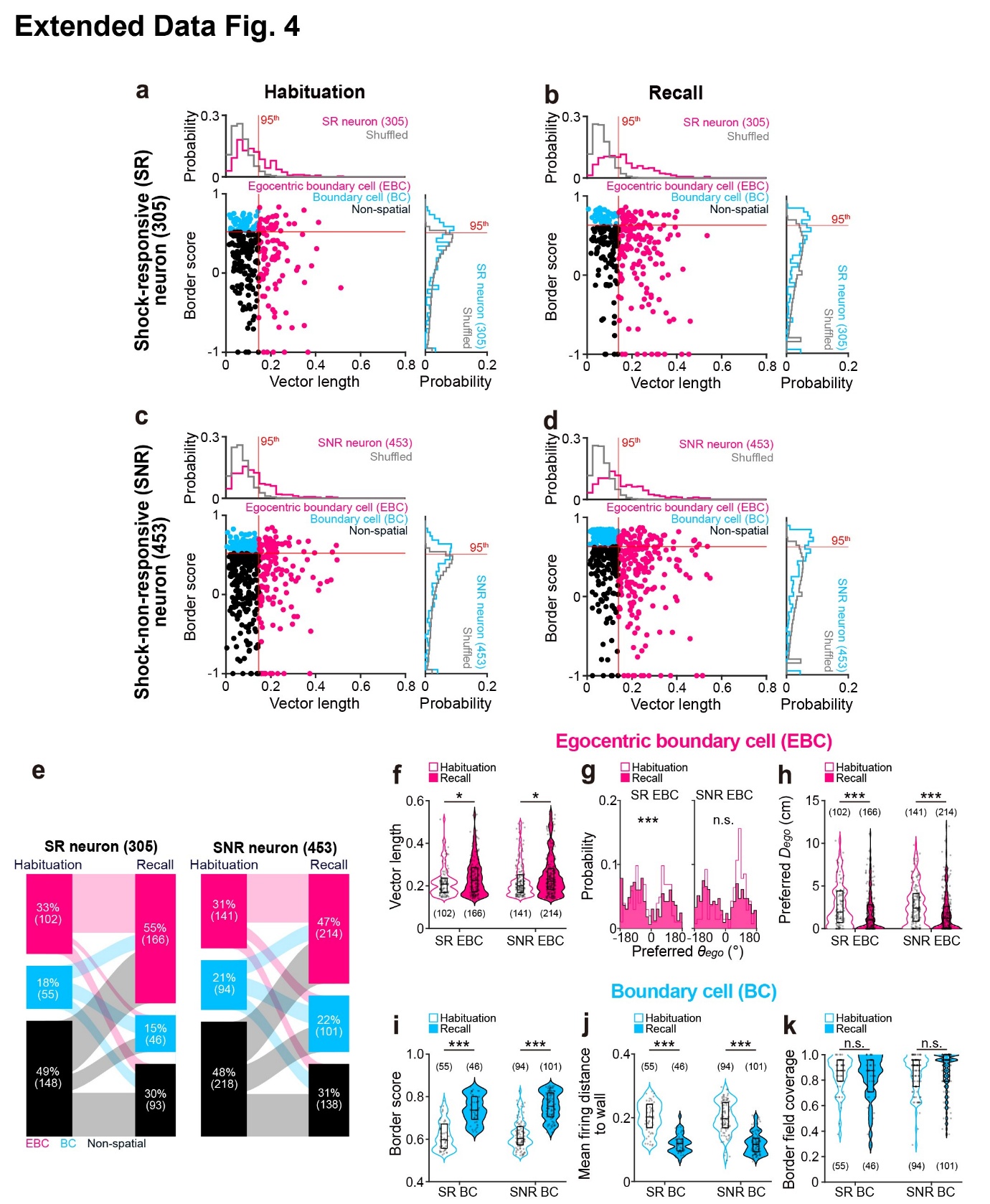

**Extended Data Fig. 4 Spatial scaffold refinements are robust to session-specific threshold choice.**

**a-b**, Scatter plot of border score as a function of egocentric vector length for shock-responsive (SR) neurons (bottom left; dots: individual neurons), with corresponding probability distributions of vector length (top left, magenta line) and border score (bottom right, cyan line) during habituation (**a**) and recall sessions (**b**). Neurons classified as egocentric boundary cells (EBCs, magenta), non-egocentric boundary cells (BCs, cyan), and non-spatial neurons (black) are indicated. Grey line: randomly shuffled distributions of vector length and border score, red line: 95^th^ percentile of randomly shuffled vector length (vertical) and border score (horizontal) distributions.

**c-d**, Same as **a-b**, but for shock-non-responsive (SNR) neurons.

**e**, Sankey plots showing the transition of EBCs (magenta), BCs (cyan), and non-spatial neurons (black) from habituation to recall sessions for SR neurons (left, n = 305 neurons in 10 mice) and SNR neurons (right, n = 453 neurons in 10 mice).

**f**, Violin plots showing the SR EBC (left) and SNR EBC vector lengths (right) during habituation (open magenta violin) and recall sessions (filled magenta violin).

**g**, Probability distribution of the preferred egocentric bearing (*θego*) of SR EBCs (left) and SNR EBCs (right) during habituation (open magenta bars) and recall sessions (fill magenta bars).

**h**, Violin plots showing the preferred *Dego* of SR EBCs (left) and SNR EBCs (right) during habituation (open magenta violin) and recall sessions (filled magenta violin).

**i-k**, Violin plots showing the border score (**i**), mean firing distance to wall (**j**), and border field coverage (**k**) for SR BCs (left) and SNR BCs (right) during habituation (open cyan violin) and recall sessions (filled cyan violin).

Box plots within violin plots indicate the 25^th^, 50^th^, and 75^th^ percentiles. Statistical comparisons were performed using Watson–Williams test (**g**), and Mann–Whitney tests (**f, h-k**). n.s. denotes not significant; * *p* < 0.05; *** *p* < 0.001. Data are presented as mean ± SEM. n = 758 neurons from 10 mice.

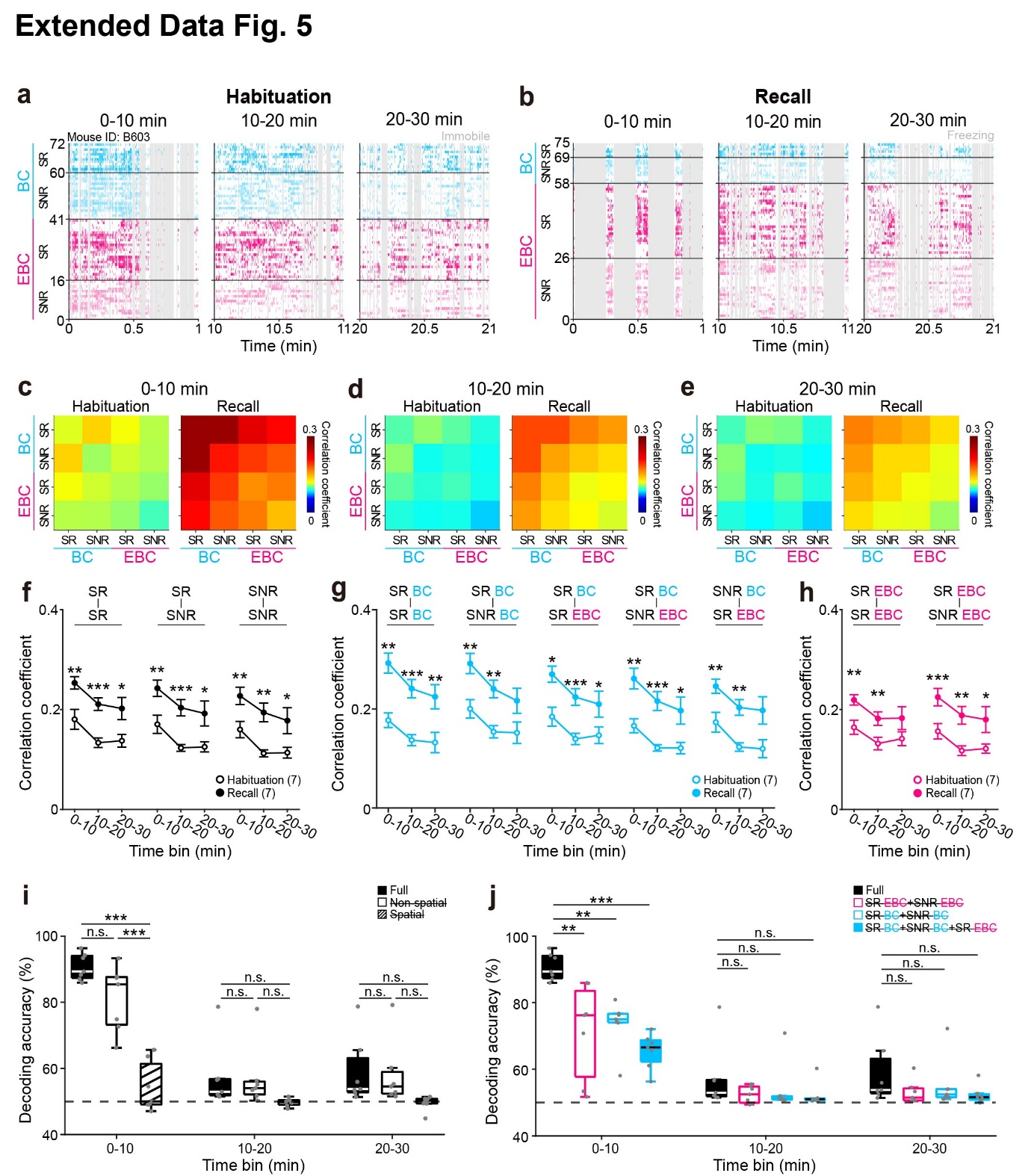

**Extended Data Fig. 5 Spatial scaffold-engram coupling is independent of locomotor speed.**

**a-b**, Representative spike rasters of shock-responsive (SR) and shock-non-responsive (SNR) egocentric boundary cells (EBCs, magenta) and non-egocentric boundary cells (BCs, cyan) across the first 1 minute of 10-min bins during habituation (**a**) and recall sessions (**b**).

**c-e**, Mean pairwise correlation matrices of SR and SNR BCs and EBCs during the 0–10 min (**c**), 10–20 min (**d**), and 20–30 min time bins (**e**) across 7 mice.

**f**, Mean pairwise correlations between SR-SR, SR-SNR, and SNR-SNR neuronal pairs across matched 10-min time bins during habituation (empty black circles) and recall sessions (filled black circles).

**g**, Mean pairwise correlations between SR BCs or SNR BCs and other spatially tuned neuronal types (SR BC, SNR BC, SR EBC, and SNR EBC) across matched 10-min time bins during habituation (empty cyan circles) and recall sessions (filled cyan circles).

**h**, Mean pairwise correlations between SR EBCs and other egocentrically tuned neurons (SR EBCs and SNR EBCs) across matched 10-min time bins during habituation (empty magenta circles) and recall sessions (filled magenta circles).

**i**, Decoding accuracy across 10-min bins following the ablation of spatial neurons (open), and non-spatial neurons (hatched), compared with all neurons (Full, filled). Grey horizontal dashed line: chance level (50%).

**j**, Decoding accuracy across 10-min bins following ablation of SR EBC + SNR EBC (open magenta), SR BC + SNR BC (open cyan), or SR BC + SNR BC + SR EBC (filled cyan) compared with all neurons (Full, filled black). Grey horizontal dashed line: chance level (50%).

Box plots indicate the 25^th^, 50^th^, and 75^th^ percentiles, and whiskers represent 1.5 times the interquartile range (**i-j**). Data are presented as mean ± SEM (**f-h**). Statistical comparisons were performed using repeated-measures ANOVA with *post hoc* Tukey’s test (**f-h**) and one-way ANOVA with *post hoc* Tukey’s test (**i-j**). n.s. denotes not significant; * *p* < 0.05, ** *p* < 0.01, *** *p* < 0.001. n = 636 neurons from 7 mice.

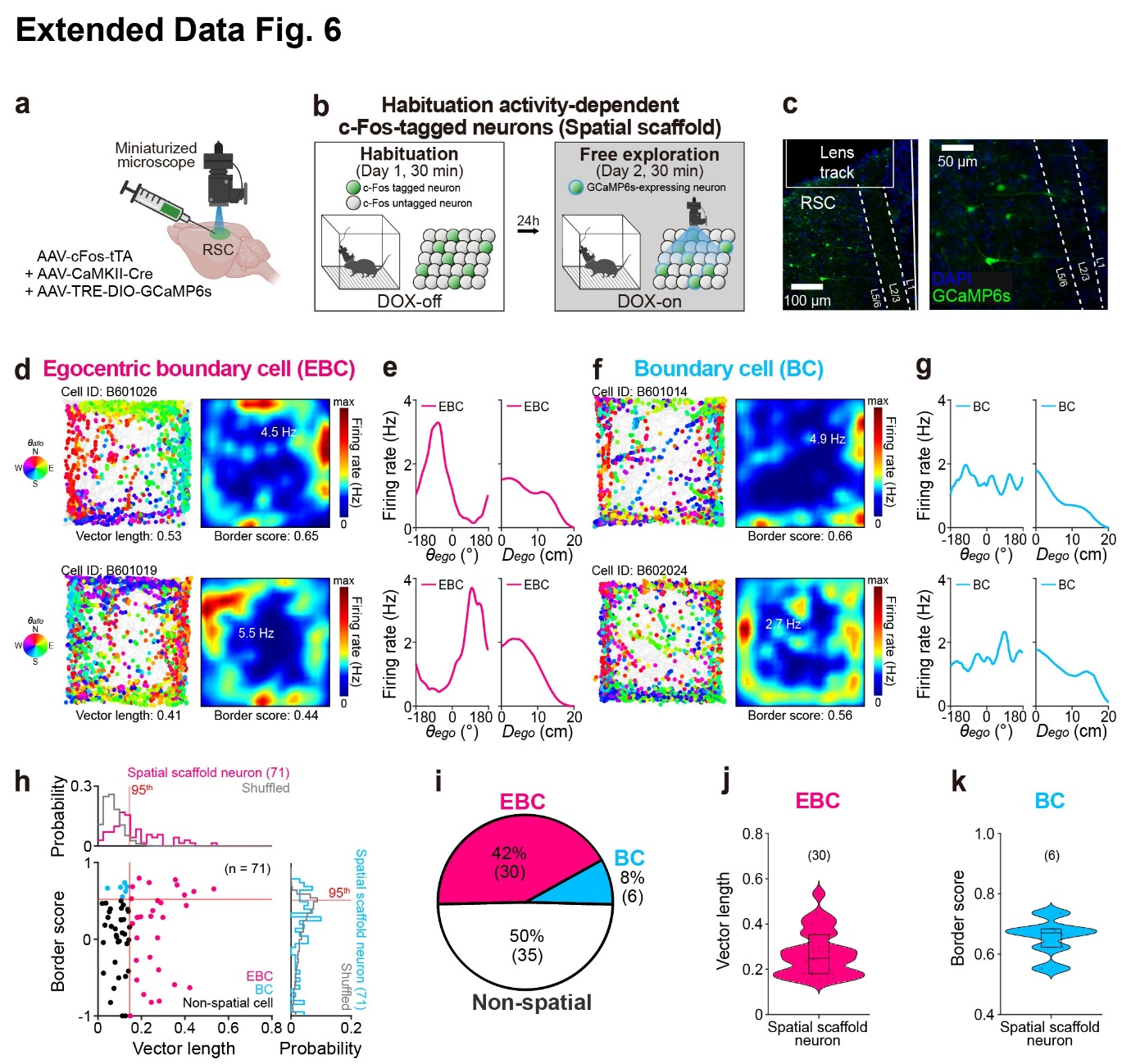

**Extended Data Fig. 6 Validation of spatial scaffold identity in habituation-tagged RSC neurons.**

**a**, Schematic of viral injection (AAV-cFos-tTA, AAV-CaMKII-Cre, AAV-TRE-DIO-GCaMP6s), GRIN lens implantation, and Ca^2+^ imaging using miniaturized microscope of c-Fos-tagged GCaMP6s-expressing excitatory neurons in retrosplenial cortex (RSC) in C57BL/6J mice.

**b**, Experimental timeline and doxycycline (DOX) schedule for habituation activity-dependent c-Fos tagging (spatial scaffold tagging). Mice underwent a 30-min habituation on Day 1 (DOX-off) for activity-dependent c-Fos-tagging, followed by 30-min free exploration on Day 2 (DOX-on) for Ca^2+^ imaging of c-Fos-tagged GCaMP6s-expressing neurons, separated by 24-h interval. Inset: Green filled circles: c-Fos-tagged neurons; Grey filled circles: c-Fos untagged neurons; Blue outline: GCaMP6s expression.

**c**, Representative fluorescent image of the GRIN lens track (left) and a magnified view (right) of c-Fos-tagged GCaMP6s-expressing neurons (green) with DAPI counterstain (blue) in RSC.

**d**, Representative spike-trajectory plots (left; mouse trajectory: grey line, spike locations: colored dots) and the corresponding spike firing rate maps (right; color scale bar: firing rate from 0 to maximum, inset: maximum firing rate, vector length, and border score) of two representative egocentric boundary cells (EBCs) recorded from c-Fos-tagged GCaMP6s-expressing RSC neurons during free exploration. Each spike is color-coded for *θallo* (inset, North: N, East: E, South: S, and West: W).

**e**, Tuning curves of egocentric bearing (*θ*ego, left) and egocentric distance (*D*ego, right) of representative EBCs shown in (**d**).

**f-g**, Same as **d-e**, but for representative non-egocentric boundary cells (BCs).

**h**, Scatter plot of border score plotted as a function of vector length for all recorded neurons (bottom left; dots: individual neurons) with corresponding probability distributions of vector length (top left, magenta line) and border score (bottom right, cyan line). Neurons classified as EBCs (magenta), BCs (cyan), and non-spatial neurons (black) are indicated. Grey line: randomly shuffled distributions of vector length and border score; Red line: 95^th^ percentile threshold of randomly shuffled distributions for vector length (vertical) and border score (horizontal).

**i**, Pie chart showing proportions of EBCs (magenta), BCs (cyan) and non-spatial neurons (white) in c-Fos-tagged RSC neuronal ensembles.

**j**, Violin plot showing the vector length of EBCs from c-Fos-tagged RSC neuron during free exploration.

**k**, Violin plot showing the border score of BCs from c-Fos-tagged RSC neuron during free exploration.

Box plots within violin plots indicate the 25^th^, 50^th^, and 75^th^ percentiles. n = 71 neurons from 2 mice.

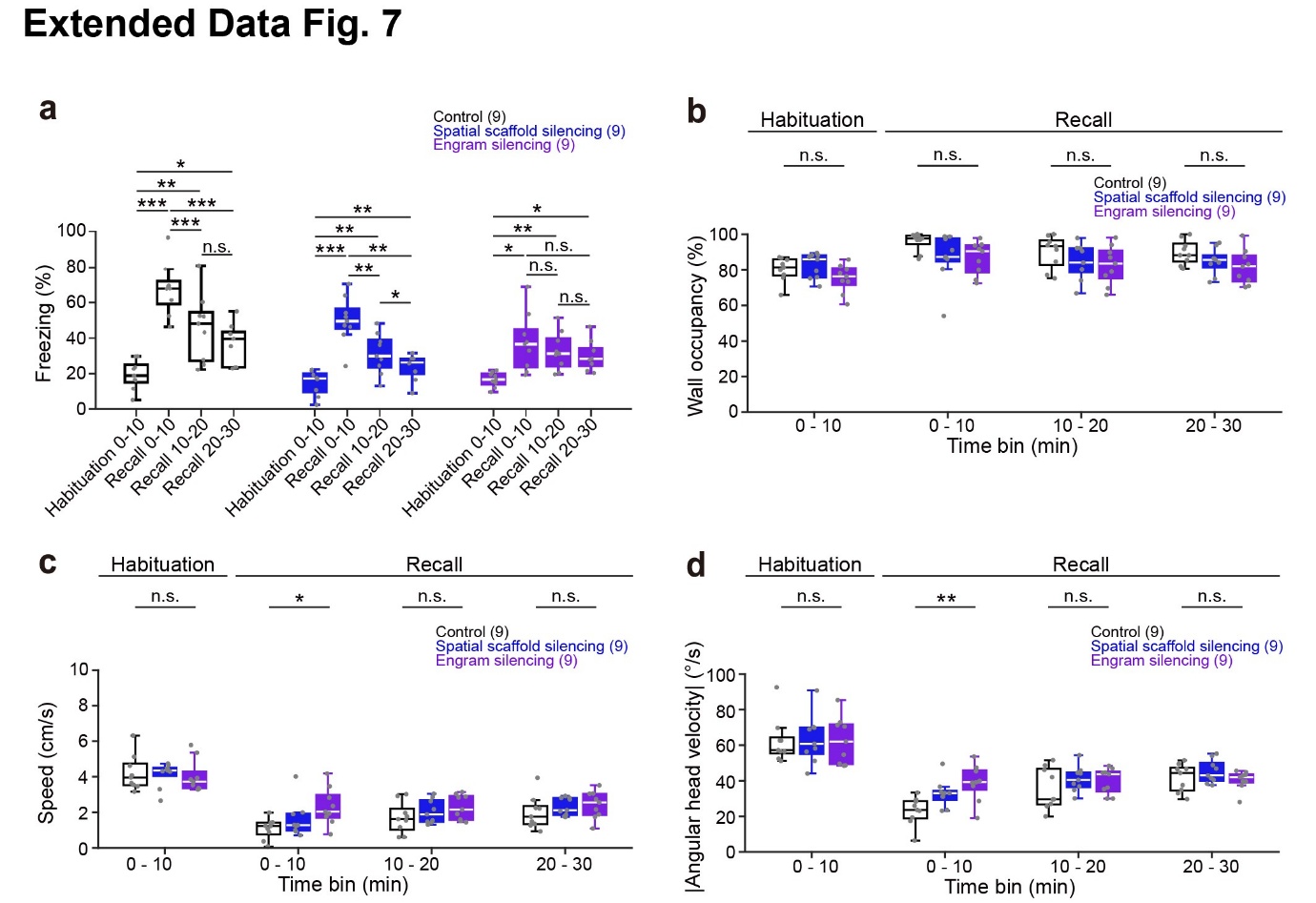

**Extended Data Fig. 7 Behavioral effects of manipulating RSC spatial scaffold and engrams during recall.**

**a**, Mean freezing during habituation (first 10 min) and recall (three 10-min bins) session in the contextual fear conditioning (CFC) paradigm in control (empty), spatial scaffold silencing (blue), and engram silencing groups (purple).

**b-d**, Wall occupancy (**b**), Mean speed (**c**), and absolute angular head velocity (**d)** during habituation (first 10 min) and the recall (three 10-min bins) session in the CFC in control (empty), spatial scaffold silencing (blue), and engram silencing groups (purple).

Box plots indicate the 25^th^, 50^th^, and 75^th^ percentiles, and whiskers represent 1.5 times the interquartile range (**a-d**). Statistical comparisons were performed using repeated-measures ANOVA with *post hoc* Tukey’s test (**a**), and one-way ANOVA with *post hoc* Tukey’s test (**b-d**). n.s. denotes not significant; * *p* < 0.05, ** *p* < 0.01, *** *p* < 0.001. n = 9 mice per group.
